# Unconventional linkers facilitate potent stabilized coronavirus stem antibody responses following nanoparticle vaccination

**DOI:** 10.1101/2025.11.19.689216

**Authors:** Christian K.O. Dzuvor, Sydney P. Moak, Lindsay R. McManus, Abigail E. Dzordzorme, Lamount R. Evanson, Taewoo Kim, Abigail Thomas, Olayimika Adeyemi, Valerie Foley, Aditi Limaye, Ryan P. McNamara, Kizzmekia S. Corbett-Helaire

**Author notes:** Authors contributed equally to this study (Christian K.O. Dzuvor, Sydney P. Moak).

## Abstract

Vaccine technologies that protect against a range of related pathogens within viral families, such as human immunodeficiency virus (HIV), influenza, and coronaviruses (CoVs) represent the future of viral vaccine development. Towards developing broad-spectrum CoV and influenza vaccines, we and others previously designed and evaluated CoV and influenza stem antigens; but these elicited relatively weak and sub-neutralizing antibody (Ab) responses. Multivalent antigen display on nanoparticles (NPs) is an established strategy to enhance and shape immunogenicity. However, one facet of NP vaccines has been largely overlooked: the indispensable linker segment between the antigen and NP core. Here, we introduce *de novo*–designed rigid (L2) and rarely used long flexible (L6) linkers to optimally display antigens on NPs, target occluded epitopes, and enhance cross-reactive Ab responses, using prefusion-stabilized Middle East respiratory syndrome coronavirus (MERS-CoV) spike (S-2P) and stem (SS) antigens as prototype antigens. Antigenic characterization of L2-NPs confirmed enhanced Ab binding and exposure of cross-reactive epitopes compared with L6-NPs and soluble antigens. Immunization with SS-L2-NPs elicited broader, more potent cross-reactive Ab responses across the seven human-infecting CoVs and pandemic threat WIV1-CoV, whereas SS-L6-NPs induced stronger neutralizing Ab responses against MERS-CoV, SARS-CoV-2, and WIV1-CoV. Ab competition and systems serology analyses revealed that SS-L2-NPs elicit robust Fc-mediated effector functions. By improving CoV-targeting Ab functionality, these linker approaches have the potential to confer broad-spectrum CoV protection and represent a promising strategy against hypervariable influenza and HIV viruses – as well as other broad viral families with pandemic potential.

## Main

Coronaviruses (CoVs) pose ongoing threats to global human health, as demonstrated by three highly pathogenic CoVs – severe acute respiratory syndrome (SARS)-CoV, Middle East respiratory syndrome (MERS)-CoV, and SARS-CoV-2 – which have emerged and re-emerged over the past two decades^1,2^. MERS-CoV, which was first identified in 2012 causing ∼1600 cases with ∼33% mortality^3^, re-emerged in Saudi Arabia in 2024 and has since caused 12 cases and at least three deaths^4^. Some recent cases could not be traced to camel reservoirs, suggesting human-to-human transmission and pandemic potential. Meanwhile, SARS-CoV-2, responsible for over seven million deaths globally, continues to circulate endemically, accumulating mutations that evade immunity.

COVID-19 vaccines, including those using rationally engineered prefusion spike (S) antigen^5^, must be updated annually to keep up with the virus’ rapid mutation rate, similar to influenza vaccines. Additional zoonotic CoVs are poised for human emergence, such as WIV1-CoV, a SARS-related virus that replicates efficiently in human cells, suggesting potential for direct transmission to humans^6,7^. Moreover, a newly identified MERS-related CoV from bats uses ACE2 receptor, the same entry receptor exploited by SARS-CoV-2^8,9^. Conservation of ACE2 usage across avian and mammalian species^10^, including humans, underscores the virus’ broad host range and potential for zoonotic spillover^11^. Before these emergences, four endemic human CoVs (HCoVs), divided into beta (HCoV-HKU1 and -OC43) and alpha (HCoV-229E and -NL63) genera, circulated seasonally causing mild respiratory infections and occasionally severe disease or death in vulnerable populations^12^. Despite alarming human health threats posed by CoVs, vaccines only exist for SARS-CoV-2.

Targeting cross-reactive epitopes – conserved regions across the CoV family – has therefore become essential for development of broad-spectrum vaccines. Such epitopes are less prone to rapid mutation, making them prime candidates for broad immune responses. Moreover, mutations in these epitopes often impose functional or structural penalties on viral fitness and survival^13,14^. CoV S mediates cellular entry and is thus the target antigen for vaccine and therapeutic development^5,15^. S comprises an S1 (head) subunit, essential for receptor engagement and viral attachment, and an S2 (stem) subunit, the spring-loaded fusion machinery. S1 contains an N-terminal domain (NTD), receptor-binding domain (RBD), and subdomains 1 and 2 (SD1 and SD2). Current CoV vaccines primarily induce humoral responses to the poorly conserved, hypervariable, and mutation-prone head region^16,17^, threatening prolonged vaccine efficacy^18^. Thus, immunodominance of S1 epitopes^19–23^ impedes induction of broad, protective, stem-directed Abs.

To advance universal CoV vaccine development^24,25^, strategies that overcome type-specific Ab responses and promote broad superfamily immunity are needed. Two complementary approaches are particularly promising^26,27^: (1) antigen design to focus on cross-reactive Ab epitopes within the immune-subdominant yet conserved S2 region^27^, and (2) nanoparticle (NP) display to enable optimal spatial organization for efficient B-cell engagement and enhanced immune responses^28^. For the former, we and others previously designed and evaluated CoV stabilized stem (SS) and influenza stem immunogens^29–33^, which lack immunodominant head domains; these vaccines elicited relatively weak Ab responses with limited cross-reactivity and bleak cross-neutralization capacity. For the latter, NPs displaying multiple CoV S proteins^21,34,35^, S1 domains^35^, or subunits (ie: RBD)^36–42^ have generally shown potentiation of immunogenicity. Many NP vaccines, including influenza and Epstein–Barr virus (EBV) NPs^43–46^, use short flexible linkers (L0) or longer flexible versions (L1) to protrude antigens from the NP core **(Fig. 1a, Supplementary Table 1)**. For example, influenza HA stem–ferritin NPs^30,44^ utilizing L0 short flexible linkers elicited non-neutralizing responses and did not provide cross-group protection. Moreover, CoV S2 antigens covalently attached to NPs via the flexible SpyTag/SpyCatcher system induced type-specific responses^47,48^ and did not elicit neutralizing Abs (nAbs)^49^.

**Fig. 1:**
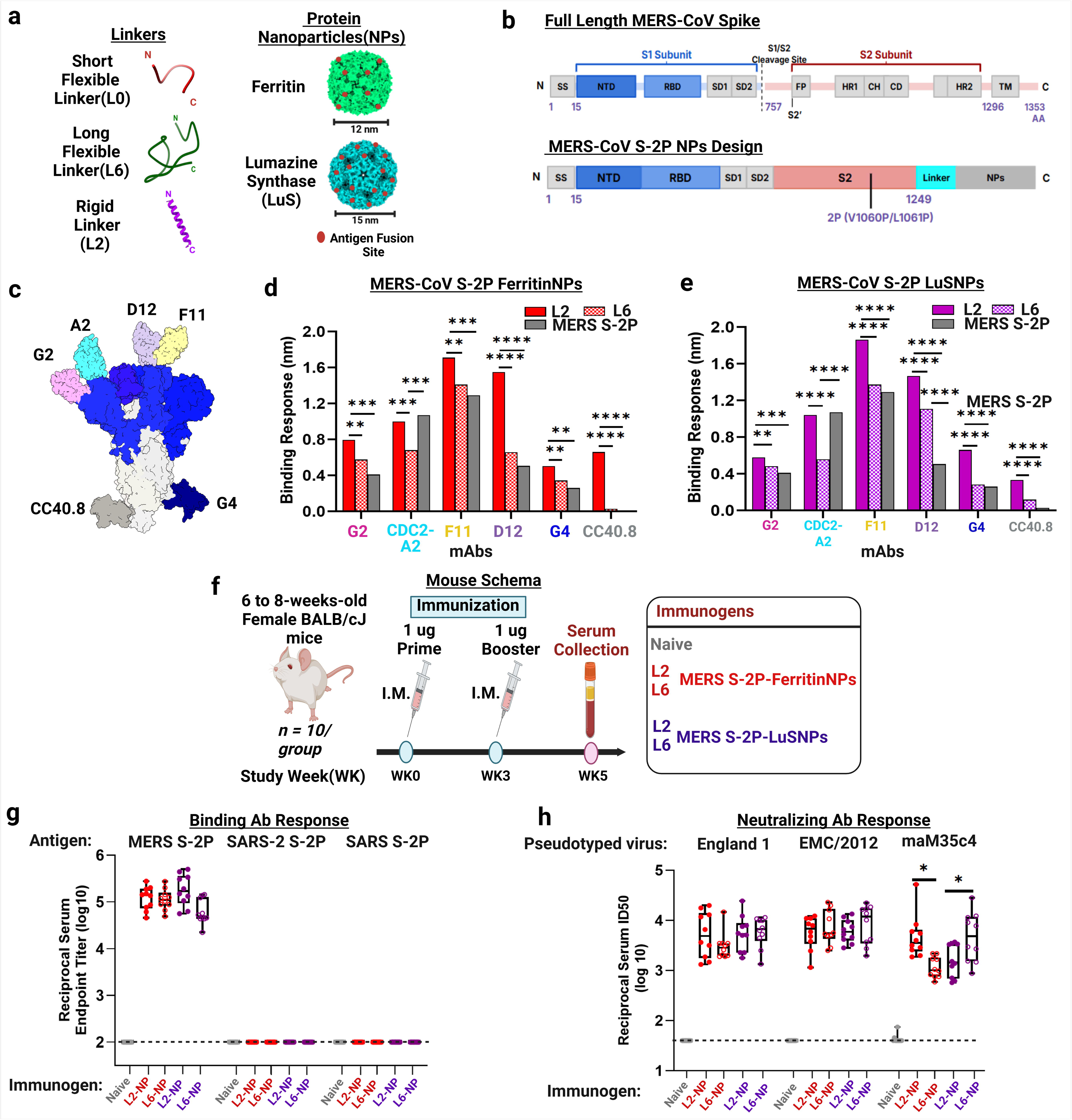
Design and immunogenicty of MERS-CoV S-2P displayed on nanoparticles using unconventional linkers (L2 and L6). (**a**) Structural models of linkers and nanoparticles (NPs). Linker secondary structures were predicted with PEP-FOLD 3. Ferritin (PDB: 1FHA) and lumazine synthase (PDB: 1HQK) were used for ferritin and LuS NPs, respectively. The antigen fusion or insertion sites are indicated as red circles. Size (nm) is indicated by reference bars. (**b**) Schematics of MERS-CoV S architecture (top) and molecular design of MERS-CoV S-2P NP construct (bottom). NTD – N-terminal domain (dark blue); RBD – receptor binding domain (light); SD – subdomain (grey); FP – fusion peptide (grey); HR1 – heptad repeat 1 (grey); CH – central helix (grey); CD – connecting domain (grey); spike neck (grey); HR2 (grey); TM (grey). S2 – stem (light red), and linker (aqua). (**c**) Model of S highlighting antibody (Ab) epitopes and monoclonal Abs (mAbs) used in biolayer interferometry (BLI): NTD (G2 - pink, A2 - aqua), RBD (D12 - purple, F11 - yellow), and S2 (CC40.8 - grey, G4 - blue). (**d,e**) Antigenic characterization of MERS-CoV S-2P-ferritinNP (**d**, red) and -LusNP (**e**, purple) by BLI. Immunogens were subjected to binding by aforementioned mAbs. Bars represent qualitative comparison of Ab binding to immunogens where L2 = solid bars, L6 = hashed bars, and soluble MERS S-2P = grey. Average of three replicates is shown. (**f**) Female BALB/cJ mice (N=10/group) were immunized at weeks 0 and 3 with 1 μg NP–Sigma Adjuvant System (SAS) adjuvant mixture and bled at week 5 for serology; immunogens used were: ferritin (red) or LuS (purple) NPs designed with L2 (closed circles) or L6 (open circles). Control(naïve) mice received adjuvanted PBS (grey). (**g-h**) Sera were assessed for binding antibody responses to MERS-CoV, SARS-CoV-2, and SARS-CoV S-2P proteins using enzyme-linked immunosorbent assay (ELISA) (**g**) and for pseudovirus neutralizing Ab responses (**h**) against MERS-CoV strains: England 1, EMC/2012, and mouse-adapted - maM35c4. Data are presented as reciprocal serum endpoint titer (log_10_) and reciprocal 50% neutralization titers, ID_50_, respectively. In the box-and-whisker plots, the horizontal line indicates median, the top and bottom of the box represent interquartile range (IQR), and the whiskers represent range. Horizontal dashed lines represent assay limits of detection (LOD). Each dot represents an individual mouse. Mouse sera with undetectable responses are overlaid on LOD. **(d-e, g-h)** Groups were compared using two-way ANOVA with Tukey’s multiple comparison test: * = p < 0.05, ** = p < 0.01, *** = p < 0.001, **** = p < 0.0001.

We reasoned that conventional flexible linkers (L0 and L1) enable moderate antigen separation but play a minimal role in protein secondary structure formation due to the absence of amino acids with side chains^50,51^. Therefore, their elicitation of limited cross-reactivity or sub-neutralizing Ab responses may result from their predominantly flexible structure, which could compromise effective B-cell receptor cross-linking^52^. Here, we present *de novo*–designed rigid helical (L2) and rarely used long flexible (L6)^53^ linkers for NP vaccine design. We hypothesize that these linkers will enable optimal orientation and display of CoV S-2P and SS on NPs, thereby targeting occluded cross-reactive epitopes and enhancing cross-reactive Ab responses. Unlike conventional flexible linkers (L0 and L1), the longer flexible (L6) linker has limited flexibility provided by amino acids with side chains such as leucine (Leu, L) and lysine (Lys, K). To evaluate effectiveness of these linkers, we utilized them to display prefusion-stabilized MERS-CoV S-2P^5^ and SS^29^ antigens on NPs.

Currently, MERS-CoV vaccines in development^54^ using S-2P elicit humoral responses primarily focused on highly variable RBD^22^, which limits breadth of protection. Similarly, display of S-2P on NPs elicits S1-focused Ab responses, while S2-targeting Abs were rarely induced^21^. Our previously designed MERS-CoV SS protein elicited sub-neutralizing responses^29^; as did SARS-CoV-2 SS protein^31,32^. Herein, improving upon immunogenicity of soluble protein, MERS-CoV S-2P–L2-NPs avidly bound the cross-reactive CoV S2-specific Ab CC40.8^55^, whereas MERS-CoV S-2P and MERS-CoV S-2P–L6-NPs did not, demonstrating that the L2 linker exposes occluded conserved or cross-reactive epitopes. Furthermore, this approach significantly improved SS cross-reactivity across seven human CoVs and pre-pandemic WIV1-CoV, while L6 linker elicited significantly higher nAb responses against MERS-CoV, SARS-CoV-2, and WIV1-CoV. Serum Ab competition and systems serology assays revealed that L2-NPs induced stronger Fc-mediated effector functions. By improving the magnitude and breadth of vaccine-induced humoral responses, these linker technologies, especially the L2 linker, impressively advance broad-spectrum NP vaccine design^28^ and may represent a novel immunofocusing strategy^26^ for hypervariable pathogens such as influenza and HIV—as well as other broad viral families with pandemic potential.

## Proof-of-concept: Exploring unconventional linkers as an immunofocusing strategy

Current MERS-CoV vaccines in pre-clinical evaluation or clinical trials^54^ using our rationally designed prefusion-stabilized S-2P antigen^5^ elicit S1-focused Ab responses. Moreover, displaying S-2P on NPs using a short flexible (L0) linker did not immunofocus such S1-targeted Ab responses towards S2^21^. S2 epitopes are occluded by S1 and are rarely targeted both in current MERS-CoV vaccine designs^21,39^ and infection^20^. Additionally, a recently isolated CoV S2-specific cross-reactive monoclonal antibody (mAb), CC40.8, was shown to bind only the conserved stem helix peptide, but not to MERS-CoV S^55^, inferring complete occlusion of these potent, cross-reactive epitope on prefusion MERS-CoV S; a similar phenomenon has been observed for some SARS-CoV-2 mAbs^56^. To that end, we sought to design NPs that could target these conserved, occluded epitopes to elicit potent, cross-reactive Ab responses.

As proof-of-concept, we first explored MERS-CoV S-2P ferritin (PDB: 1FHA) and lumazine synthase (LuS)(PDB: 1HQK) NPs, using two unconventional linkers (L2 and L6) (**Fig. 1a–b, Supplementary Table 1**). The MERS-CoV S-2P (1–1294 amino acid) ectodomain maintains 2P substitutions and S1/S2 furin cleavage site mutations (RSVR-to-ASVG), as originally described^5^; we include deletion of 47 C-terminal S ectodomain residues guided by prior SARS-CoV-2 SΔC-ferritin design^57^. We hypothesized that truncating S-2P’s ectodomain would promote robust expression and optimal NP display, thereby enhancing stability and immunogenicity. MERS-CoV S-2P fused to foldon trimerization domain^58^, designed in our previous study^5^, was used as a benchmark. Compared with MERS-CoV S-2P, NPs showed intended size and architecture by size-exclusion chromatography (SEC), dynamic light scattering (DLS), and negative-stain transmission electron microscopy (NS-TEM) (**Extended Data Fig. 1a–d**). As expected, differential scanning fluorimetry (DSF) revealed that NPs possess higher melting temperatures compared with MERS-CoV S-2P (**Extended Data Fig. 1e–f**), suggesting that linker display NPs did not alter S-2P stability.

We next evaluated antigenicity of these NPs by biolayer interferometry (BLI) using a panel of mAbs specific to NTD (G2^59^, CDC2-A2^60^), RBD (F11^59^, D12^59^), and S2 (G4^59^, CC40.8^55^) domains (**Fig. 1c**). MERS-CoV S-2P, S-2P-L2- and -L6-NPs revealed significant differences in Ab binding (**Fig. 1d-e**) for both ferritin and LuS NPs. Both L2-NP and L6-NP bound strongly to CDC2-A2 than MERS-CoV S-2P. L2-NPs’ enhanced binding to all mAbs compared to L6-NPs and MERS-CoV S-2P, suggesting more optimal display of MERS-CoV S-2P on NPs using L2 linker. Interestingly, L2-NPs avidly bound the cross-reactive stem helix-specific mAb CC40.8^55^, whereas L6-NPs and S-2P did not, indicating that the L2 linker display approach has potential to expose transmembrane-proximal occluded cross-reactive epitopes.

Finally, we evaluated immunogenicity of these MERS-CoV S-2P NPs by measuring humoral binding Ab and nAb titers. Groups of 10 six to eight-weeks-old BALB/cJ mice were immunized intramuscularly (IM) at weeks 0 and 3 with Sigma Adjuvant System (SAS)–adjuvanted NPs (**Fig. 1f).** Two weeks post-boost, both L2- and L6-NP elicited ∼5-log MERS-CoV S-2P specific reciprocal endpoint binding titers; however, no cross-reactive Ab responses to SARS-CoV or SARS-CoV-2 S-2P were detected (**Fig. 1g**). Our findings are consistent with previous studies^21,39^ for MERS-CoV S-2P-L0-I53-50NP. This outcome may be due to elicited Ab focused on immunodominant S1^5,20^, which is not well conserved across CoV clades (**Extended Data Fig. 2a–c**). Nonetheless, sera from NP-immunized mice elicited robust, broad MERS-CoV cross-clade nAbs, with median 50% neutralization titers (ID_50_) ranging from ∼3 - 4 logs (**Fig. 1h**). L2-NP elicited significantly higher nAb titers against mouse-adapted MERS-CoV (maM35c4) than L6-NP for ferritin and vice versa for LuSNP. Collectively, these data demonstrate production of stable, antigenically intact, and highly immunogenic MERS-CoV S-2P NPs, using distinct linkers. Moreover, these data suggest our de novo-designed L2 linker supersedes L6 in its ability to target an occluded S2 epitope.

## Design, *in vitro* assembly, and characterization of SS–nanoparticle immunogens

Since MERS-CoV S-2P NPs failed to elicit cross-clade reactive Ab responses, we focused efforts toward using L2- and L6-NPs to improve cross-reactivity and overall immunogenicity of MERS-CoV S2, the more cross-reactive S domain (**Extended Data Fig. 2b–c**). Targeting S2 is one of the universal CoV vaccine field’s goals, as highlighted by previous design and evaluation of CoV SS immunogens^24,25^. However, as yet, CoV SS elicits robust cross-reactive Ab responses with little to no neutralization capacity; complete cross-clade CoV protection has not been achieved, or at least not tested^29,31–33^.

To produce MERS-CoV SS NPs, we genetically fused SS to three NP backbones^28^: (1) ferritin, (2) LuS, and (3) I53-50 using L2 and L6 linkers, which represent a breadth of size and antigen-particular density (**Fig. 2a–b**). Ferritin, LuS and I53-50 NPs display eight, twenty and sixty trimeric antigens, respectively^28^. Analytical SEC purification of expressed proteins revealed well-defined peaks corresponding to NP and SS protein immunogens, indicating efficient assembly and formation (**Extended Data Fig. 3a-b**). Comparison of SS-NPs to SS and bare NPs by DLS and NS-TEM revealed monodisperse, homogeneous, well-defined spherical NP assemblies with apparent additional densities on the NP surface (**Extended Data Fig. 3c–d**).

**Fig. 2:**
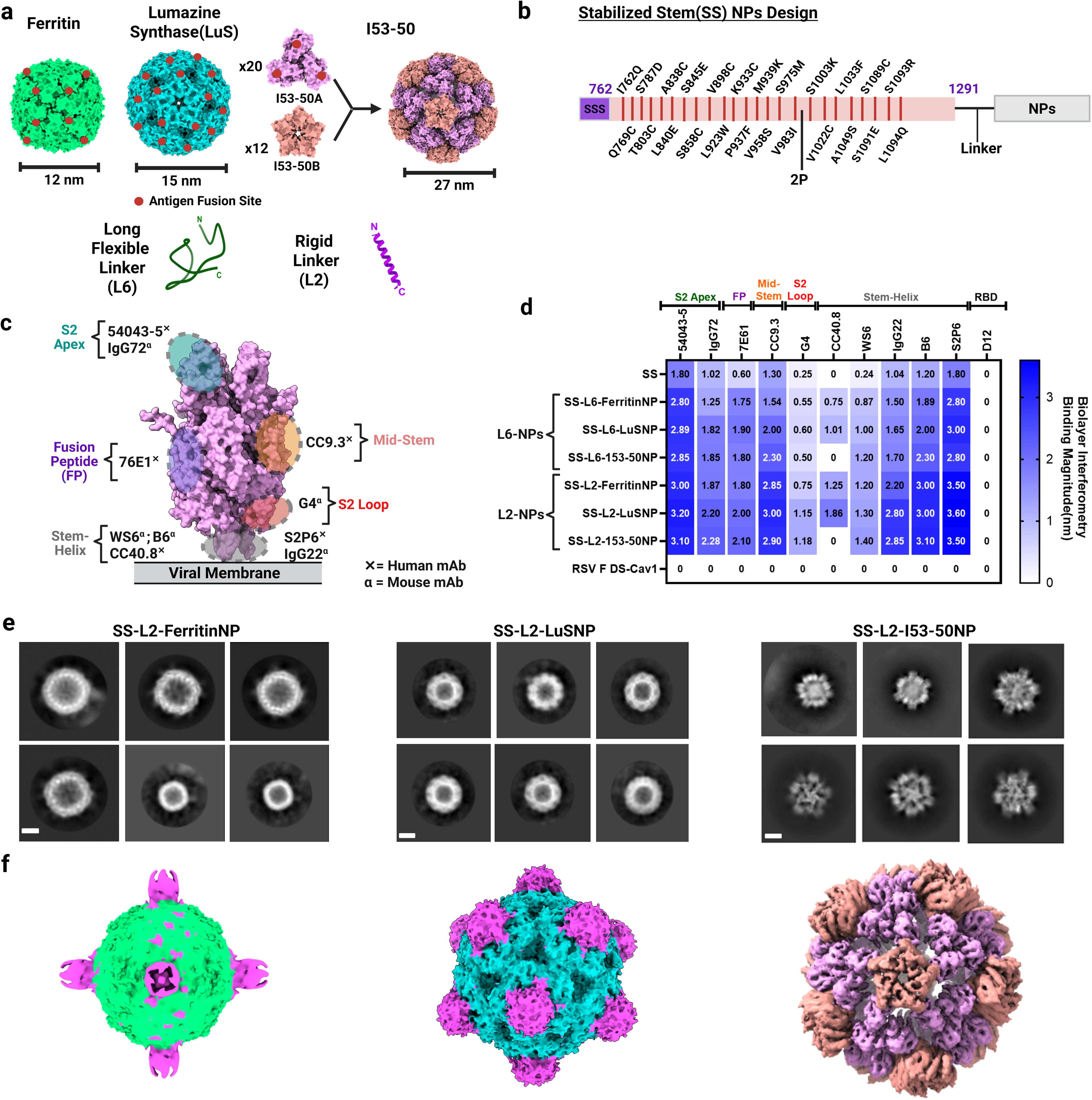
Design and characterization of MERS-CoV SS-NPs. (**a**) Structural models of ferritin (PDB: 1FHA), LuS (PDB: 1HQK), and I53-50 (PDB: 7SGE) NPs and linkers. Engineered fusion sites for antigen display are indicated as red dots. (**b**) Schematic representation of stabilized S2 stem (SS) fused to NPs via linker. Stabilizing mutations are shown by red lines. Secretion signal sequence (SSS) is shown in purple. (**c**) Epitopes targeted by cross-reactive S2-specifc mAbs used in BLI. (**d**) Antigenic characterization of immunogens bound to S2 mAbs as assesed by BLI, where binding strength is measured in nm and represented by a color gradient: dark blue (strong) to white (no binding). (**e**) Two-dimensional (2D) class averages of cryo-electron microscopy (EM) micrographs of MERS-CoV SS-ferritin (left), -LuS (middle), and -I53-50 (right) NP. Scale bar, 20 nm. (**f**) Single-particle cryo-EM reconstruction of SS-L2-NPs, determined at a resolution of 6.25, 4.07, and 4.42 Å for ferritin, LuS, and I53-50 NPs, respectively. Cryo-EM density maps were fitted with atomic models (in Fig. 2a) and colored accordingly.

Next, we assessed the immunogens’ antigenicity against a panel of cross-reactive S2-specific mAbs (**Fig. 2c**). All NPs showed robust binding profiles compared with SS, with L2-NPs exhibiting the highest Ab binding (**Fig. 2d**). Additionally, we imaged L2-NPs by cryo-electron microscopy (cryo-EM). Single-particle cryo-EM reconstructions of cryo-frozen NPs were obtained at 6.25, 4.07, and 4.42 Å resolutions for ferritin, LuS, and I53-50 NPs, respectively (**Fig. 2e–f; Extended Data Figs. 4–6; Supplementary Table 2**). While SS was visible in ferritin and LuS NPs, it was poorly resolved for I53-50 NPs after particle image averaging and data processing (**Extended Data Fig. 5**). Nonetheless, fitting NP models into the density map supported design precision and demonstrated that genetic fusion of SS to NPs via our novel L2 linker did not alter SS or NP architecture (**Fig. 2f**). Together, these data show production of antigenically intact SS–NPs containing unconventional linkers.

## Stability of SS–nanoparticle immunogens

To determine the immunogens’ resilience and ability to meet stringent stability requirements of manufacturing, storage, and distribution^61^, we monitored SS and NP stability via DSF and storage at −80°C,4°C, 37°C, 25°C, and over 30-days, using SDS–PAGE and SEC analyses. As expected, DSF analysis showed that NPs manifested higher or similar thermal stability compared with SS alone (**Extended Data Fig. 3e–g**), suggesting that linker display on different NP backbones did not have deleterious effects on SS folding. As well, SS-L2-LuSNP were resistant to degradation at −80°C,4°C, and 25°C, up to 30 days (**Extended Data Fig. 7a, upper**). Incubation at 37°C demonstrated exceptional stability, with partial degradation appearing on days 14 and 30. In contrast, SS degraded severely at 4°C, 25°C, and 37°C after 4 days of incubation (**Extended Data Fig. 7a, lower)**. After 30 days, SEC analysis revealed that NPs remained more stable at 4°C, 25°C, and 37°C compared with SS, as observed by well-defined and regular chromatograms (**Extended Data Fig. 7b**). Under similar conditions, SS chromatograms appeared irregular, indicating protein degradation or aggregation (**Extended Data Fig. 7b)**.

We further assessed retention of protein architecture via epitope-specific mAb binding after thermal stress. Compared with SS, SS-L2-LuSNP showed no discernible loss in antigenicity (**Extended Data Fig. 7c**). SS-L2-LuSNP bound mAbs IgG72^29^ and G4^59^ equivalently after incubation for 1 h at different temperatures, suggesting that L2-NP display further stabilizes the prefusion conformation of SS, perhaps due to L2’s rigidity and NP backbone. Collectively, these data demonstrate that NP immunogens exhibit superior thermal and storage stability, above that of soluble MERS SS protein.

## Elicitation of cross-reactive antibody responses

To evaluate cross-reactivity, we immunized BALB/cJ mice IM with 10 µg of MERS-CoV SS as soluble trimers or displayed on L2 and L6 NPs formulated with SAS adjuvant. Adjuvanted phosphate-buffered saline (PBS) served as a negative control. We immunized mice at weeks 0 and 3 and collected blood at week 5 to assess serum Ab responses (**Fig. 3a**). MERS-CoV SS protein and both NPs elicited similar levels of homotypic MERS-CoV S-2P-binding Ab responses at ∼5 logs median reciprocal EC_50_ titers (**Fig. 3b**), as expected;10 µg of SS alone was previously shown to max out MERS-CoV-specific binding Ab responses^29^. Interestingly, L2 significantly outperformed L6 in ferritin and/or I53-50 NPs, eliciting more robust SARS-CoV-2, SARS-CoV, WIV1-CoV, and HCoV-HKU1 S-2P–specific Ab responses, with reciprocal median EC_50_ titers in the range of 4.5 to 5.5 logs. However, with LuSNP backbone, L2- and L6-NPs performed equally, eliciting, for example, SARS-CoV-specific Ab responses at ∼4.5 reciprocal median EC_50_ titer. Generally, both NPs elicited significantly more robust Ab responses than soluble SS (**Fig. 3a-f**). Soluble SS failed to elicit HCoV-OC43 S-2P–specific Ab responses, as previously reported^29^. Notably, displaying SS on NPs rescued SS’s ability to elicit HCoV-OC43–specific Ab responses, with L2-ferritin NPs significantly outperforming L6-ferritin NPs – a testament to L2-NPs’ ability to enhance cross-reactivity (**Fig. 3g**). These results suggest that NP display of MERS-CoV SS overall induces greater cross-reactivity than soluble SS. L2 linker display of SS generally elicits more cross-reactivity than the L6 linker; except for LuSNP, which may allude to density-dependent effects on L2 linker’s ability to immunofocus on cross-reactive S2 epitopes.

**Fig. 3:**
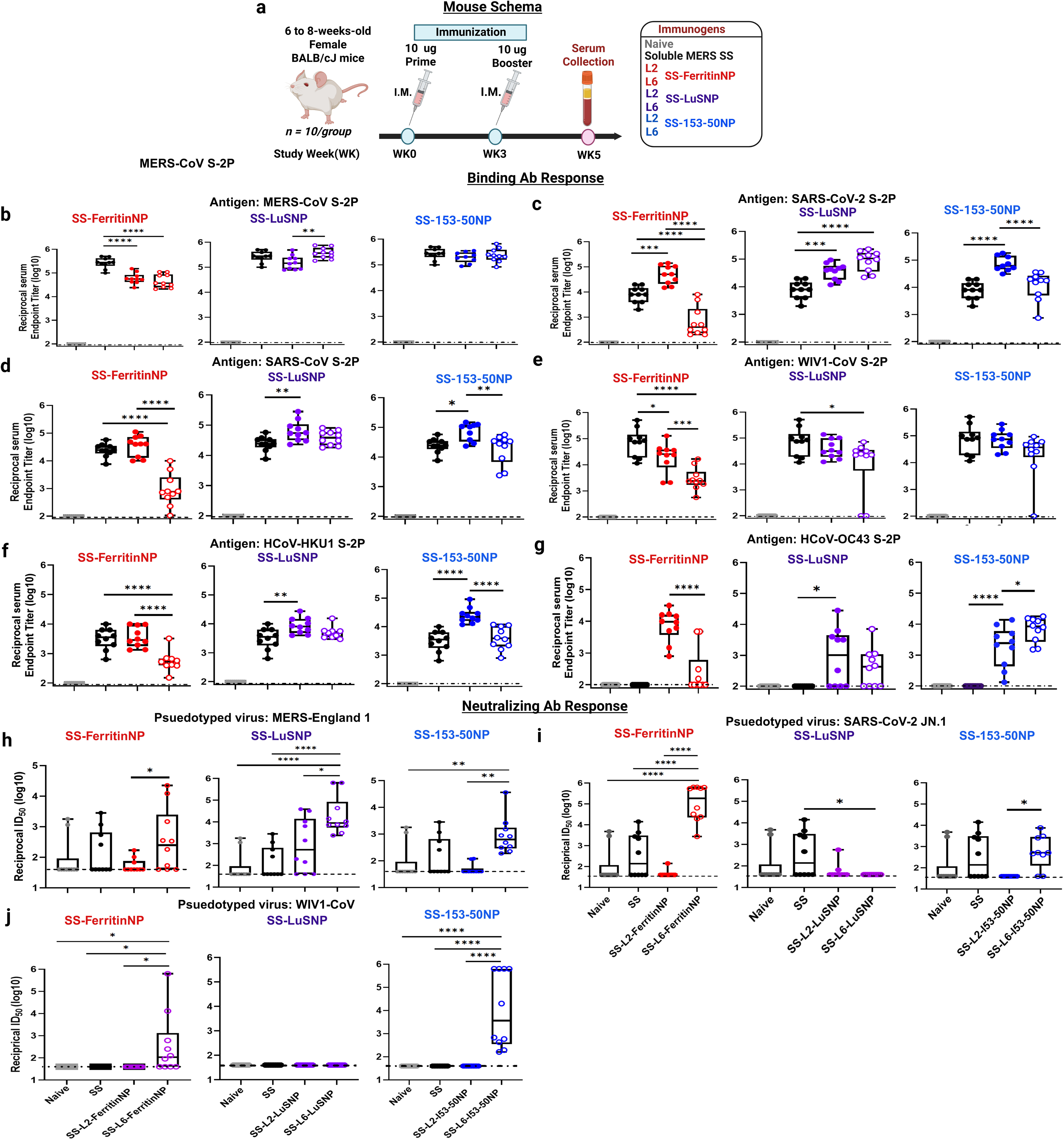
Immunogenicity of MERS-CoV SS-NP vaccines in mice. (**a**) Female BALB/cJ mice (N=10/group) were immunized at weeks 0 and 3 with 10 μg NP–SAS adjuvant mixture and bled at week 5 for serology. Mice were immunized with MERS-CoV SS ferritin (red), LuS (purple), or I53-50 (blue) NPs designed with L2 (closed circles) or L6 (open circles) linkers. Negative and historic control mice received adjuvanted PBS (grey) or soluble SS (black). (**b–j**) Sera were assessed for binding Ab responses to S-2P proteins from MERS-CoV (**b**), SARS-CoV-2 (**c**), SARS-CoV (**d**), WIV1-CoV (**e**), HCoV-HKU1 (**f**), and HCoV-OC43 (g) using ELISA and for pseudovirus neutralizing responses against MERS-CoV (h), SARS-CoV-2 (**i**), and WIV1-CoV (**j**). Data are presented as reciprocal serum endpoint titer (log_10_) and reciprocal 50% neutralization ID_50_ titer, respectively. In the box-and-whisker plots, the horizontal line indicates median, the top and bottom of the box represent IQR, and the whiskers represent range. Horizontal dashed lines represent assay limits of detection (LOD). Each dot represents an individual mouse. Mouse sera with undetectable responses are overlaid on LOD. Groups were compared using two-way ANOVA with Tukey’s multiple comparison test: * = p < 0.05, ** = p < 0.01, *** = p < 0.001, **** = p < 0.0001.

## Elicitation of autologous and heterologous neutralizing antibody responses

Given NPs elicited robust cross-reactive binding Ab responses, we next assessed pseudovirus neutralizing activity. As previously reported^29^, soluble SS elicited only modest homotypic MERS-CoV nAb responses, with only 4 out of 10 (40%) SS-immunized mice having detectable responses (>2 logs ID_50_). Interestingly, L6 linker facilitated significantly stronger MERS-CoV nAb responses than L2 for all three NP backbones, with median ID_50_ values ranging from ∼2 to 3 logs, suggesting that L6 effectively focuses Abs responses on type-specific MERS-CoV S2 Ab epitopes **(Fig. 3h)**. This result was not limited to homotypic responses; SS-L6-NPs induced significantly more robust SARS-CoV-2 and WIV1-CoV nAb responses than SS-L2-NPs, for ferritin and I53-50 backbones. SS-L2-LuSNP did not elicit detectable SARS-CoV-2 or WIV1-CoV nAb responses (GMT ID_50_ < 2 logs) (**Fig. 3i–j**). These results suggest that the cross-reactive Ab responses elicited by SS-L2-NPs have orthogonal, non-neutralizing functions.

## Epitope specificity of polyclonal antibody responses

To dissect Ab epitope specificity in the polyclonal immune mouse sera towards pinpointing additional Ab functionalities, we carried out competition BLI^19^ **(Supplementary Fig. 1)** competing the mouse sera with well-characterized S2-specific mAbs (**Fig. 2c**). We used mAbs recognizing five major antigenic sites: S2 apex (54043-5^62^), S2 fusion peptide (76E1^63^), S2 mid-stem (CC9.3^64^), S2 stem helix (CV3-25^65^, S2P6^66^, B6^67^, IgG22^29^), and S2 loop (G4^59^) to investigate a breadth of cross-binding and cross-neutralizing epitopes (**Supplementary Fig. 1a**). By way of brief background: stem apex–targeting Abs, like 54043-5, are highly cross-reactive but non-neutralizing^62^; fusion peptide–targeting Abs, such as 76E1, are highly cross-reactive and potently neutralizing^63^; stem helix– and mid-stem–targeting Abs display modest cross-reactivity^64,68–70^; S2 loop–targeting G4 Ab is MERS-CoV-specific^59^ and dominates Ab responses following MERS-CoV SS vaccination^29^.

Anti-Penta-HIS (HIS1K) biosensors coated with SS trimers were first incubated with serially diluted immune sera and then with one of the mAbs. As serum concentration increased during primary binding, secondary Ab binding declined (**Supplementary Fig. 1b**), indicating vaccination produced Abs that bound to epitopes recognized by these mAbs in a dose-dependent and specific manner. Percentage competition was calculated as compared to previous studies^19,46,71^. Ferritin and Lus L2-NPs outperformed SS and L6-NPs, by eliciting the most abundant stem apex-targeting 54043-5-like Abs (∼88-96%) (**Fig. 4a-c**). Notably, although these Abs cross-react with all seven human CoVs (**Supplementary Fig. 1a**), they lack neutralizing activity. 54043-5-like S2 apex Abs confer cross-protection through Fc-mediated effector functionality^62^. Fc-mediated mechanisms have also been proposed as protective modes of action for clinical influenza HA stem-based immunogens^30,43,44,72^, suggesting a conserved mechanism of protection across viral families. In contrast, SS elicited more mid-stem-targeting CC9.3-like Abs (77%) than both L2- and L6-NPs (∼27-65%). Nonetheless, the most cross-nAb class, 76E1, was also strongly elicited by Ferritin and LuS L2-NPs (∼41- 53%) over SS and L6-NPs (∼7-12%) **(Fig. 4a-c)**. Moreover, SS-L6-FerritinNP elicited more CV3-25-like Ab (70%), while all other stem-helix-targeting Abs were approximately equally elicited; interestingly, SS-L6-FerritinNP did not elicit as many 54043-5-like Abs (only ∼39%) as the other SS-NPs **(Fig. 4a).** Together, this pattern may indicate immunodominance of the 54043-5 epitope in L2-NPs.

**Fig. 4:**
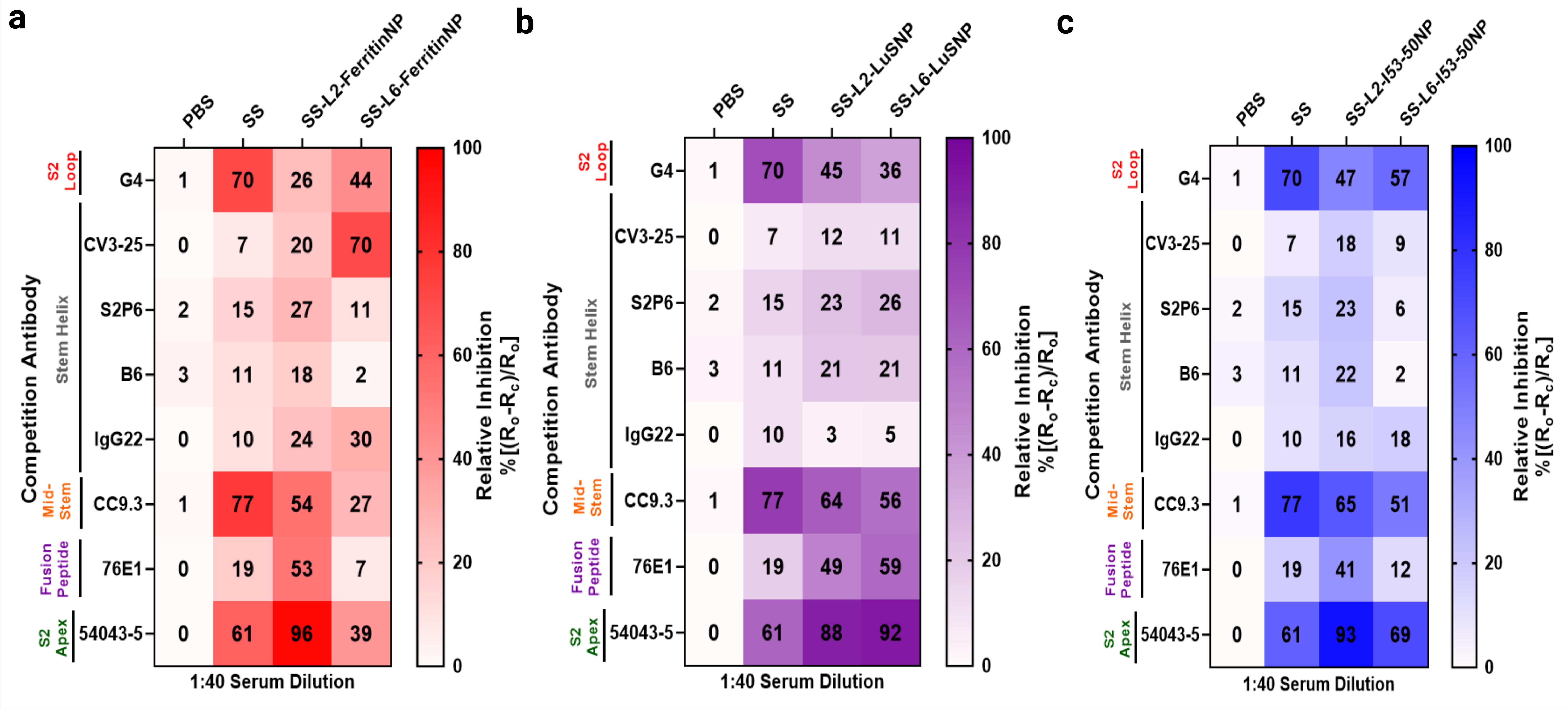
**Dissecting epitope specificity of polyclonal Ab responses in MERS-CoV SS-NP-immune mouse serum. (a-c**) Heatmaps of the % of relative competition of pooled mouse sera with mAbs (see Fig. 2c for mAbs description and epitopes). Mice were immunized with ferritin (**a**, red), LuS (**b**, purple), and I53-50 (**c**, blue) SS-NPs. Results from (Ro − Rc) × 100 / Ro represent competition levels, where Ro is the highest binding signal of mAb without serum and Rc is that with serum from vaccinated mice. Dilution or inhibition level is represented as a color gradient, where strongest inhibition is darkest and weakest inhibition is lightest.

As expected, MERS-CoV soluble SS elicited ∼70% Abs that competed with mAb G4, aligning with our prior findings that MERS-CoV elicits robust G4-like Abs^29,59^. Here, Strikingly, MERS-CoV–specific G4-like Ab responses were lower in both L2-NPs (∼26-47%) and L6-NPs (∼36-57%) groups than in SS-vaccinated mice, indicating that both L2 and L6 linkers immunofocus Ab responses away from this MERS-CoV-specific epitope. This is promising as one goal of our unconventional linker NP display approach is to decrease type-specificity while increasing cross-reactivity.

Together, these results suggest that SS-L2-NPs may elicit overpowering amounts of Fc-mediated effector functioning Abs, such as 54043-5-like Abs, and perhaps Abs target novel cross-reactive but sub-neutralizing epitopes. Interestingly, these data also infer that the robust cross-neutralization elicited by SS-L6-NPs is likely attributed to a yet-discovered class of cross-nAbs.

## Stimulation of Fc-mediated antibody effector mechanisms

Given the low levels of nAb responses and the substantial amount of cross-reactive 54043-5-like^62^ Abs elicited, we next investigated alternative Ab-mediated mechanisms of action. We profiled polyclonal sera from individual immunized mice using systems serology assays (**Supplementary Fig. 2**)^73^. First, we determined binding Abs to several stabilized S and SS proteins (MERS-CoV S-2P, SARS-CoV-2 S-2P, HCoV-HKU1 S-2P, HCoV-OC43 S-2P, SARS-CoV S-2P, WIV1-CoV S-2P, MERS-CoV SS, SARS-CoV SS), subclass specificity (Total IgG, IgG1, IgG2a, IgG3, IgM, IgA), and interactions with specific Fcγ receptors (FcγR IIb, FcγR III, or FcγR IV). Ebolavirus glycoprotein (EBOV GP) was used as a negative response control. We used naïve sera (adjuvanted PBS group) as a binding reference control (**Fig. 5a, Supplementary Fig. 2a**). As observed in our ELISA assay (**Fig. 3, Extended Data Fig. 8a**), immune sera contained high levels of IgG, IgG1, IgG2a, IgG3, and IgM Abs against S and SS proteins but not EBOV GP compared with naïve sera (Fig**. 5a**). Moreover, immune sera showed higher binding to both inhibitory (FcγR IIb) and activating FcγRs (FcγR III and FcγR IV) for most S proteins except EBOV GP–coated control beads (**Fig. 5a**). No binding nor FcγR-binding responses were observed for EBOV GP for any group, as expected (**Fig. 5a**).

**Fig. 5:**
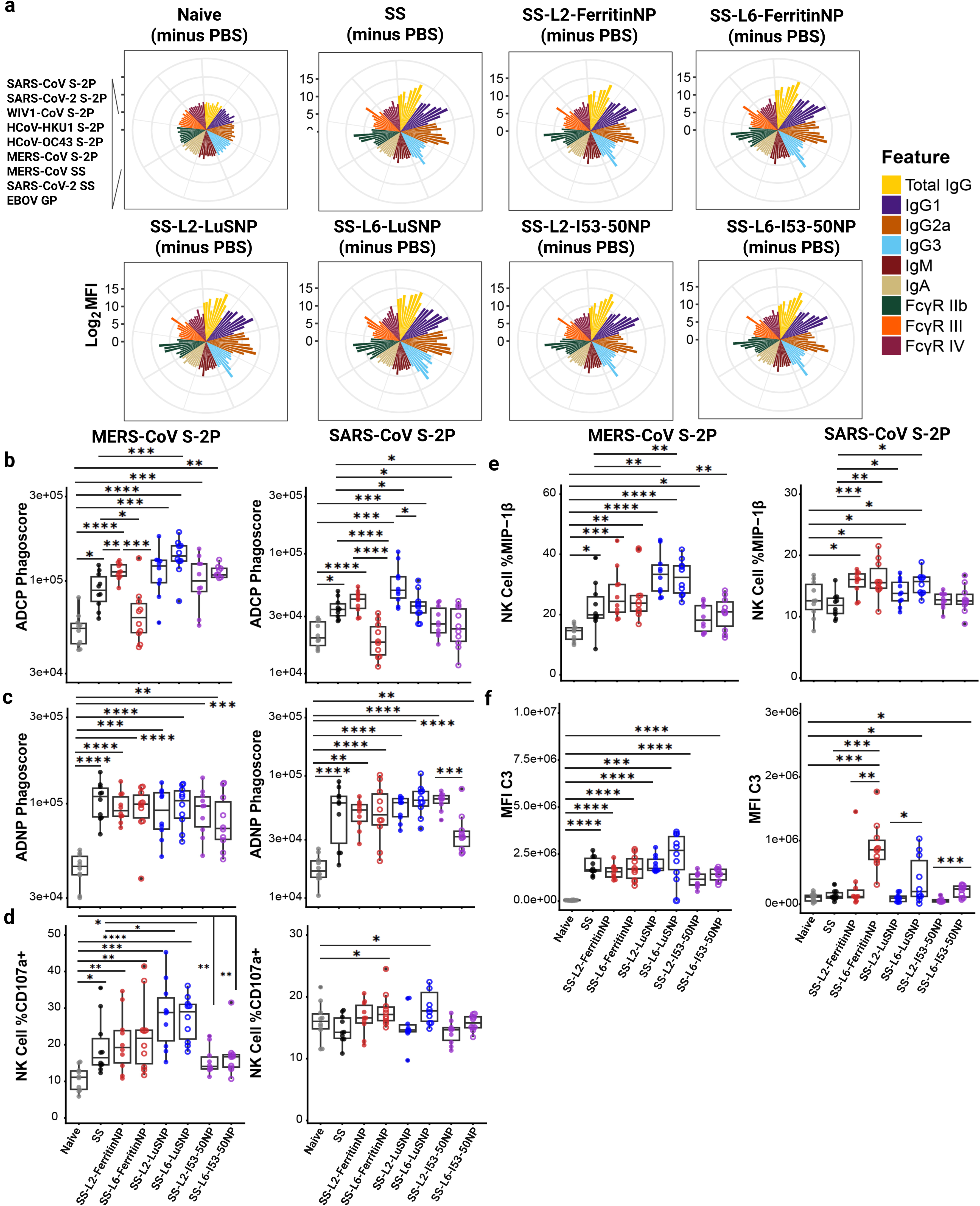
Systems serology analysis of MERS-CoV SS-NP vaccine-induced immune sera. (**a**) Radar plots representing biophysical analysis of Ab and FcγR-binding titers of naïve and immune sera after immunization with soluble MERS-CoV SS and SS-NPs. All binding features were background (PBS) subtracted and plotted in log2 binding MFI. Shown on the right is the color scheme for the binding features. On the left is the antigen order. (**b–f**) Ab Fc-mediated effector functions of naïve and immune sera against MERS- and SARS-CoV S-2P proteins. Ab-mediated cellular phagocytosis with monocytes (**b**, ADCP) or neutrophils (**c**, ADNP) using vaccine-induced immune and naïve sera and beads coated with MERS-CoV and SARS-CoV S-2P proteins. (**d–e**) Ab-mediated natural killer (NK) cell activation (ADNKA) using naïve and vaccine-induced immune sera and beads coated with MERS-CoV and SARS-CoV S-2P proteins as quantified by (**d**) percentages of NK cells surface with surface expression of CD107a and (**e**) percentages of NK cells expressing MIP-1β. For all cellular assays, whole blood or leukopacks from two independent donors were used. (**f**) Ab-dependent complement deposition (ADCD) on beads coated with MERS and SARS-CoV S-2P proteins and incubated with naïve or immune sera. For all box-and-whisker plots, the horizontal line indicates median, the top and bottom of the box represent interquartile ranges, and the whiskers represent the range. Each dot represents an individual mouse. Groups were compared using two-way ANOVA with Tukey’s post hoc test: *p < 0.05; **p < 0.01; ***p < 0.001; ****p < 0.0001.

Given the enhanced FcγR-binding profiles of immune sera from vaccinated groups, we next assessed Fc-effector functions, focusing on Ab-dependent cellular phagocytosis by monocytes (ADCP), Ab-dependent neutrophil phagocytosis (ADNP), Ab-dependent NK cell activation (ADNKA), and Ab-dependent complement deposition (ADCD). Each assay was tested against MERS-CoV and SARS-CoV S-2P proteins to evaluate the breadth of Fc-effector responses (**Fig. 5b–f, Supplementary Fig. 2b-d**). ADCP activity was induced by most vaccine groups with the exception of SS-L6-Ferritin NP to MERS-CoV S-2P. Likewise, SS-L6-ferritin NP did not show significant ADCP to SARS-CoV S-2P either. All other groups showed strong ADCP to MERS-CoV S-2P (**Fig. 5b**, left). Interestingly, a trend was observed for SS-L2 vs. SS-L6 NP vaccine groups for cross-reactive ADCP to SARS-CoV S-2P (**Fig. 5b**, right). For ADNP, all groups had a statistically significant response to MERS-CoV S-2P compared to naive, including SS and NP treatment arms (**Fig 5c**, left). For cross-reactive ADNP responses to SARS-CoV S-2P, we observed strong responses for all groups except for SS-L6-I53-50 NP. Again, when contrasted to the L2 counterpart, the L6 showed reduced effector function (**Fig. 5c**, right).

For ADNKA analysis, we measured the percentages of NK cells expressing CD107a, a degranulation marker, and macrophage inflammatory protein 1 beta (MIP-1β), an NK cell activation marker and pro-inflammatory cytokine. Compared with naïve sera, immune sera significantly enhanced ADNKA, as reflected by increased CD107a (**Fig. 5d)** and MIP-1β (**Fig. 5e**) expression to MERS-CoV S-2P. Percentages of CD107a+ and MIP-1β+ NK cells were markedly higher in both SS-L2-LuSNP and SS-L6-LuSNP groups compared with the SS group. Vaccination with SS-L6-ferritinNP and SS-L6-LuSNP significantly increased the percentage of CD107a^+^ NK cells compared with all other groups in response to SARS-CoV S-2P. In contrast, all vaccinated animals except those in the SS and SS-I53-50NP groups exhibited significantly higher MIP-1β+ NK cell percentages in response to SARS-CoV S-2P,compared to naïve (**Fig. 5e**), with SS-L2-FerritinNP and SS-L6-LuSNP groups seemingly exhibiting the strongest effects. ADCD, as quantified by C3 deposition, was significantly induced for immune sera from all groups in response to MERS-CoV S-2P compared with naïve sera (**Fig. 5f**). Immune sera from SS-L6-NP–vaccinated mice promoted ADCD against SARS-CoV S-2P.

Overall, these findings demonstrate that vaccination with either construct elicited diverse cross-reactive Abs with Fc-mediated effector mechanisms of protection. The Linkers (L2 vs. L6) showed effector-specific responses, mostly for SARS-CoV S-2P, with L2-NPs producing the most pronounced effects for opsinophagocytosis and L6-NPs showing enhanced complement fixing.

## Model of epitope exposure, possible B-cell activation, and immune induction

Our work uncovers unconventional linkers that enable optimal display of MERS-CoV S-2P and SS antigens on NPs, allowing for targeting of cross-reactive – and sometimes neutralizing – Ab epitopes. Based on antigenic and functional characterizations, we propose B-cells stimulated by SS-L6-NPs result in Ab responses that are limited in cross-reactivity and reduced cross-reactive Fc-effector functions, but impressively neutralizing, particularly for Ferritin- and LuS. On the other hand, and importantly as we aim towards increased Ab breadth with promiscuous protective functionality, L2’s rigid helical orientation on SS-NPs likely facilitates strong B-cell receptor cross-linking and activation, leading to potent cross-reactive Abs with Fc-mediated effector functionality and potentially other orthogonal functions, such as cell-to-cell fusion blockade^74^ (**Fig. 6**).

**Fig. 6:**
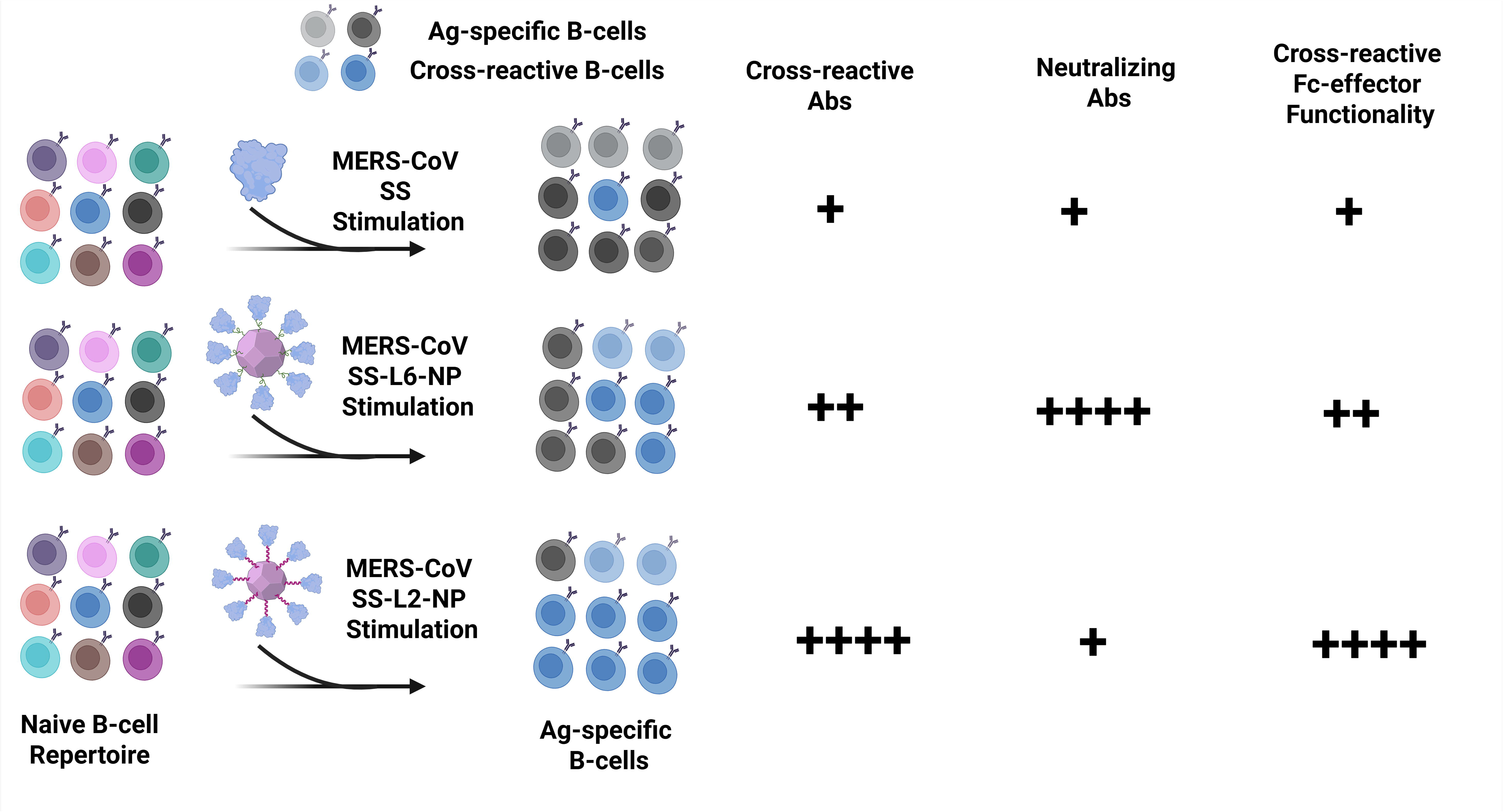
Proposed model of immune response induction inferred from antigenicity, immunogenicity, and profiling of polyclonal sera from immunized mice. Model of the ability of MERS-CoV soluble SS (top), L6- (middle), and L2-NPs (bottom) to stimulate B-cells towards induction of cross-reactive Abs, neutralizing Abs, and cross-reactive Fc-effector functions. Antigen-specific and cross-reactive B-cells are colored grey and blue, respectively.

## Conclusions

Vaccines eliciting broadly cross-reactive immunity against current human-infecting CoVs and diverse CoVs that are poised for emergence are urgently needed^25^. Current CoV vaccines^24,27^ induce Ab responses primarily directed towards immunodominant epitopes in spike S1 subunit, which are poorly conserved and prone to escape mutations^60,75,76^, thereby threatening durable vaccine efficacy. Towards pandemic preparedness, it is imperative to develop broadly protective vaccines. To that end, immunofocusing on subdominant conserved S2 epitopes provides a promising avenue.

Proof-of-concept immunofocusing strategies (as recently reviewed)^26,77^, such as protein dissection, epitope scaffolding, epitope masking, cross-strain boosting, and conventional multivalent NP display, elicit subfamily-specific Ab responses with weak and limited functionality. For example, CoV SS immunogens^29,31–33^ were designed to focus Ab responses on conserved stem supersites; however, responses elicited were relatively weak, sub-neutralizing, and not cross-protective. Multivalent CoV S protein or S subunit NPs, typically using L0 and L1 linkers, elicit higher immune responses, albeit with limited viral clade cross-reactivity or neutralization^35,37,39,41,47–49,57^. Specifically, Merbecovirus S2 NPs, using SpyTag/SpyCatcher^78^, did not elicit nAbs^49^. Recently, non-covalent display using an engineered nanobody enabled modular projection of multiple viral antigens onto NPs for broad-spectrum immunization^79^. However, this approach can induce undesirable anti-nanobody Abs, which can dampen cross-reactive responses, and nanobody production using alpacas is expensive and labor-intensive^80^.

Therefore, we envision optimal antigen spacing and projection on NPs using orthogonal more rigid linkers (L2 and L6) will immunofocus on cross-reactive Ab epitopes to confer broad superfamily immunity. In this study, we explored a *de novo*–designed rigid L2 and rarely used long flexible, but modestly rigid, L6 linkers to display MERS-CoV S-2P and SS on NPs. In all, the NPs targeted occluded epitopes and elicited cross-reactive binding, neutralizing, and Fc-mediated effector functioning Ab responses. Specifically, MERS-CoV S-2P-L2-NPs enhanced Ab binding and exposure of a cross-reactive epitope, CC40.8. SS-L2-NPs elicited broad cross-reactive Ab responses across the seven human CoVs and pre-pandemic WIV1-CoV. Furthermore, SS-L2-NP-elicited Ab responses targeting diverse cross-reactive S2 epitopes: S2 apex (54043-5^62^), fusion peptide (76E1^63^), mid-stem (CC9.3^64^), and stem helix (CV3-25^65^, S2P6^66^, B6^67^, IgG22^29^) - all the while reducing SS’s accessibility of MERS-CoV type-specific epitope, G4^59^. Notably, SS-L6-NPs elicited strong nAb responses against MERS-CoV, SARS-CoV-2, and WIV1-CoV, which requires further experimental elucidation. While Ab responses for SS-L6-FerritinNP was pinpointed to be dominated by broadly nAb, CV3-25^65^, hypothetically, SS-L6-NPs may be eliciting novel Abs that recognize uncharacterized cryptic epitopes with signature motifs common across beta-CoVs, such as recently identified nAb, 3D1^81^. In addition to neutralization, we observed that NP immunogens could stimulate several Ab effector functions that were cross-reactive. Previous work has shown that these non-neutralizing functions leveraged by FcγR-binding Abs protect against antigenically drifted or mis-matched targets^82,83^. L2-LuS NP’s inability to outperform L6-LuSNP may be attributed to density-dependent effects on L2 linker’s ability to immunofocus on cross-reactive and neutralizing S2 epitopes, and needs to be probed further.

Antigen’s spatial organization, repetition and spacing on virions have long been argued to play key roles in immune responses to infection^84,85^. Prior studies have demonstrated the importance of these parameters in vaccination, suggesting that an optimal distance of 5–15 nm may be immunogenic ^86–88^. However, applying this knowledge to improve vaccine design has been difficult due to the paucity of tools to produce systematic series of protein NPs immunogens with varied features of high precision. Using our linker technologies, we emphasize the importance of optimal antigen spacing, patterning and extension from NPs^28^, corroborating others’ findings^37,89–91^. Indeed, antigen organization, patterning and spatial arrangement are cornerstones of NP vaccines’ immunogenicity^92^. To the best of our knowledge, only a handful of approaches have elicited such cross-reactive responses, including mosaic RBD^40,89^, S^21,34,93^, and immunodominant domain (IDD)^19^ NPs. Nonetheless, NPs displaying singular heterologous RBDs showed no substantial difference in overall immunogenicity^37,89^. Displaying homologous SS on NPs via our linkers renders a much broader response across the CoV family. Thinking ahead, NP display of a singular antigen lends to more reliable engineering and manufacturability.

Finally, although investigating the protective efficacy of these NP immunogens is vital, the strong cross-reactive and functional Ab responses elicited by these NPs are predictive of protection^94^. Also, evaluating how NPs’ immunization influence T-cell responses, particularly via their mRNA-encoded versions^53,95,96^, is needed to assess the full potential of these linker technologies. It will also be critical to tie our Ab results to B-cell functionality and memory. Because our results point to ability of these NPs to elicit yet-discovered broadly cross-reactive and/or neutralizing Abs, isolating novel pan-CoV mAbs from immunized mice will uncover more about CoV S2 antigenic landscape and perhaps lead to pan-CoV mAb therapies. Furthermore, future studies will need to systematically evaluate additional linkers^50,97^ (particularly, longer flexible with extra amino acids with side chains and rigid/semi-rigid with proline residues) for more in-depth understanding of linker-type specific effects on NPs’ immunogenicity, an endeavor that will be behooved by multiplex screening and powerful machine learning methods.

In conclusion, developing universal vaccines has progressed to sophisticated NP display with promising enough results to have propelled candidates into clinical trials^98–103^. However, NPs containing conventional flexible linkers have limited ability to elicit broad, cross-clade immunity and neutralization. Our data suggest that the “missing link” to attaining desirable broad multi-functional Ab responses may be use of unconventional linkers, particularly our *de novo*-designed L2 linker, to protrude antigens from NP cores. Beyond CoVs, L2-NPs can be used to display antigens from other hypervariable viruses, such as HIV or influenza, providing promise towards next-generation vaccines for pandemic preparedness.

## Methods

### EXPERIMENTAL MODEL AND SUBJECT DETAILS

#### Cell lines

HEK293T/17 (CRL-11268, ATCC) 293T cell lines, which stably overexpress human ACE2 (MF_293T-hACE2^104^; supplied by M. Farzan, Scripps Research Institute), were maintained in Dulbecco’s modified Eagle’s medium (DMEM) supplemented with 10% FBS, 2 mM glutamine, and 1% penicillin-streptomycin. For MF_293T-hACE2^104^, 3 µg/mL puromycin was added. HUH 7.5 cells (kindly provided by Robert Roth, Apath L.L.C) were cultured in DMEM supplemented with 10% FBS, 2mM glutamine, and 1% penicillin-streptomycin. All monolayer cell lines were cultured at 37°C with 5% CO_2_ and passaged to maintain a density of 1–1.3 × 10^6^ cells/mL. Cells were regularly tested for mycoplasma contamination and remained negative. Suspension Expi293F cells (#A14527, ThermoFisher), were cultured and maintained in Expi293 expression medium at 37°C, 8% CO_2_, and 70% humidity, shaking at 200 rpm. Cells were passaged and maintained twice weekly at 2–3 × 10^5^ viable cells/mL.

#### Mice

Mouse experiments were conducted in accordance with the guidelines, regulations, and policies of the National Institutes of Health (NIH) for the Care and Use of Laboratory Animals. Protocols were approved by the Institutional Animal Care and Use Committee at Harvard University and the Center for Comparative Medicine at Harvard T.H. Chan School of Public Health. Six- to eight-weeks-old female BALB/cJ mice (#000651, Jackson Laboratory) were housed in groups of five under a 12 h light/12 h dark photoperiod, at an ambient room temperature (RT) of 18–22°C and a relative humidity of 50 ± 5%. Mice were fed standard chow diets. Blood collection was performed via terminal cardiac puncture under anesthesia, and every effort was made to minimize animal suffering during this procedure.

### METHOD DETAILS

#### Construction of expression plasmids

All NPs constructs were human codon-optimized, synthesized, and cloned into the pcDNA3.1(+) mammalian expression vector by GenScript for protein expression in Expi293F cells. Both SS and S-2P NPs consisted of previously designed MERS-CoV SS-V2^29^ and MERS-CoV S-2P^5^ genetically fused to the N-terminus of ferritin, LuS, or the trimeric I53-50A.1NT1 NP component^28^ using the following linkers: L2 (GGGGSAEAAAKEAAAKEAAAKAGGGG), L6 (GGGSGGGGSGGGGSGLSK) (**Supplementary Table 3)**

For BLI assays, previously designed plasmids encoding heavy and light chains of mAbs [(G2, D12, F11, G4)^59^, (CDC2-A2)^60^, WS6^105^], [(IgG72, IgG22)^29^ and 54043-5^62^], (B6)^67^ and (CC40.8,CC9.3)^64^ were provided by Barney Graham (NIH, National Institute of Allergy and Infectious Disease, Vaccine Research Center – NIH, NIAID, VRC), Jason McLellan (University of Texas, Austin), David Veesler (University of Washington), and Raiees Andrabi (Scripps Research Institute), respectively. For mAb, S2P6^66^, purified protein was provided by Humabs Biomed SA (Bellinzona, Switzerland). Plasmids expressing heavy and light chains of mAbs (76E1^63^, FP.006^106^, CV3-25^65^) were synthesized and cloned into the pcDNA3.1(+) mammalian expression vector by GenScript.

For S glycoproteins used in ELISA, multiplex antigen binding, and systems serology assays, codon-optimized sequences of MERS-CoV (GenBank accession AFY13307.1), SARS-CoV (GenBank accession AAP13441.1), SARS-CoV-2 (GenBank accession QHD43416.1), WIV1-CoV (GenBank accession KC881007), HCoV-HKU1 (GenBank accession ABC70719.1), and HCoV-OC43 (GenBank accession AIL49484.1) S-2P^5^, previously reported in this study^21^, were obtained from Barney Graham (NIH, NIAID, VRC).

Plasmids expressing MERS-CoV and SARS-CoV-2 SS^29,32^ were obtained from Jason McLellan (University of Texas, Austin). All S-2P and SS constructs contained a C-terminal foldon trimerization motif^58^ (YIPEAPRDGQAYVRKDGEWVLLSTFL), an octa-histidine tag (HHHHHHHH), and a twin Strep-tag II (WSHPQFEKGGGSGGGSGGGSWSHPQFEK) for purification. The protein constructs included a human rhinovirus 3C protease recognition site (GSRSLEVLFQGP) for tag cleavage after purification. For lentiviral plasmids, codon-optimized full-length S sequences of MERS-CoV (GenBank accession, AFY13307.1), SARS-CoV-2 (GenBank accession QHD43416.1), and WIV1-CoV (GenBank accession KC881007); pHR’ CMV Luc (luciferase reporter gene); and pCMV DR8.2 (lentivirus backbone), cloned into the pCMV vector as previously described^21^, were provided by Barney Graham (NIH, NIAID, VRC).

## Computational and sequence analysis

For phylogenetic analysis, sequences of the seven HCoV S protein domains (S1 and S2) were aligned using Clustal Omega^107^. Phylogenetic trees were constructed using default settings in DNASTAR software (Madison, WI, USA), and sequence identity between S1 and S2 domains of the different CoVs was determined. The aligned complete S sequences were used to develop a multiple sequence alignment (MSA), which was plotted and overlaid onto MERS-CoV S (Protein Data Bank, PDB ID: 5W9J), chain A. The resultant S chain A was recolored based on conservation and reproduced to replace S chains B and C. The data were visualized using UCSF Chimera X^108^. The sequences used for generating the MSA were SARS-CoV-2 (GenBank accession QHR63290.2), SARS-CoV-1 (GenBank accession AAP41037.1), MERS-CoV (GenBank accession ALK80242.1), HCoV-HKU1 (GenBank accession ADN03339.1), HCoV-OC43 (GenBank accession QEG03814.1), HCoV-229E (GenBank accession ABB90529.1), and HCoV-NL63 (GenBank accession QED88026.1).

## Immunogen and spike expression

Spike and nanoparticle immunogens were produced by transient transfection of the respective expression vectors using polyethyleneimine “PEI-MAX” (#24765, Polysciences) as previously described with minor modifications^5^. Briefly, Expi293F cells grown in Corning polycarbonate Erlenmeyer flasks, shaking at 37°C, 8% CO_2,_ 80% humidity, and 120 rpm to a density of 3 × 10^6^ cells/mL, were transfected at a 1:3 (v/v) ratio consisting of 1 mg plasmid to 1 mg/mL PEI-MAX in Opti-MEM Reduced Serum Medium. Next, an enhancer, HyClone SFM4HEK293 medium (#SH30521.02, Cytiva), was added 16–18 h post-transfection. Cells were harvested 5–6 days post-transfection via centrifugation at 4000 × g for 30 min. Culture supernatants containing secreted proteins were filtered with a 0.45 μm filter (#4612, Pall Corporation). The respective secreted proteins were purified as described below.

## Immunogen and spike purification

### Ni-NTA affinity chromatography

Stabilized SS, S-2P, and SS-I53-50A.1NT1 constructs were purified using Ni-NTA affinity chromatography via their octa-histidine tags. Filtered supernatants were diluted and incubated overnight with Ni-NTA agarose beads (#30210, Qiagen) at 4°C to allow protein binding. The beads were then washed with 10 column volumes (CV) of wash buffer (1× PBS, 50 mM imidazole). Bound proteins were eluted with 20 CV of elution buffer (1× PBS, 300 mM imidazole) and concentrated using a 30 kDa cut-off centrifugal device (#MAP001C37, Pall Corporation). The proteins were further purified by size-exclusion chromatography using a Superose 6 Increase 10/300 GL column (#28990944, Cytiva) on an ÄKTA Pure system (Cytiva, Marlborough, MA, USA) in running buffer (1× PBS, pH 7.4). Protein purity was determined by SDS–PAGE. Prior to immunization, octa-histidine tags for SS and SS-I53-50A.1NT1 proteins were cleaved using HRV-3C protease (#Z03092-100, GenScript) following the manufacturer’s protocol. Protein concentrations were determined with a NanoDrop Lite spectrophotometer (ThermoFisher, Waltham, MA, USA) using the protein’s molecular weight and extinction coefficient obtained from the ExPASy ProtParam tool (https://web.expasy.org/protparam/).

### GNA lectin affinity chromatography

SS–ferritin NPs and SS–LuS NPs were purified via *Galanthus nivalis* lectin affinity chromatography. Briefly, 5 mL *Galanthus nivalis* lectin agarose beads (#AL-1243-5, Vector Laboratories) were combined with clarified supernatants and incubated overnight at 4°C. Next, NP–agarose suspensions were loaded onto a disposable column and washed with 20 CV of 1× PBS (pH 7.4) to remove unbound proteins and impurities. Bound NPs were eluted with 1 M methyl-α-D-mannopyranoside (#M6882, Sigma) in 1× PBS buffer (pH 7.4). NPs were further purified by size-exclusion chromatography using a Superose 6 Increase 10/300 GL column on an ÄKTA Pure system. NP purity was assessed by SDS–PAGE. Protein concentrations were determined with a NanoDrop Lite spectrophotometer (ThermoFisher, Waltham, MA, USA) using molecular weight and extinction coefficient values calculated with the ExPASy ProtParam tool (https://web.expasy.org/protparam/).

## Antibody expression and purification

mAbs were produced in Expi293F cells by transient co-transfection of heavy chain and light chain plasmids in a 1:1 ratio using polyethyleneimine (PEI-MAX). After 5 days post-transfection, cell supernatants were harvested, filtered with a 0.2 µm filter, and purified with MabSelect PrismA agarose beads (#17549801, Cytiva) following standard procedures. The beads were extensively washed with 1× PBS (pH 7.4), and Abs were eluted with 0.1 M glycine (pH 2.9) into one-tenth the volume of neutralization buffer (1 M Tris-HCl, pH 8.0). The purified Abs were concentrated, buffer-exchanged into PBS using a 30 kDa cut-off centrifugal device, and flash-frozen for long-term storage at −80°C. Ab concentrations were determined using a NanoDrop Lite spectrophotometer (ThermoFisher, Waltham, MA, USA). Abs were validated for binding by BLI before long-term storage and use.

## I53-50B.4PT1 expression and purification

The I53-50B pentamer was expressed as previously described^109^ in Lemo21 (DE3) cells (#C2528J, NEB). Cells transformed with I53-50B.4PT1 were grown in LB medium (10 g tryptone, 5 g yeast extract, 10 g NaCl) at 37°C to an OD600 of 0.7 in a 2 L shaker flask. The temperature was then reduced to 18°C, protein expression was induced with 1 mM IPTG, and cells were grown for an additional 16 h. Cells were harvested and lysed by ice sonication (amplitude 50%, time 5 min, pulse duration 15 s ON/45 s OFF) in lysis buffer (50 mM Tris, 500 mM NaCl, 30 mM imidazole, 1 mM PMSF, 0.75% CHAPS). Lysates were clarified by centrifugation at 20,000 × g for 30 min and incubated with Ni-NTA agarose beads for 1 h for purification. Protein-bound Ni-NTA agarose beads were washed with ten column volumes of buffer (50 mM Tris, 500 mM NaCl, 30 mM imidazole, 1 mM PMSF, 0.75% CHAPS), and protein was eluted with buffer containing 300 mM imidazole. The eluates were pooled, concentrated using a 10 kDa cut-off centrifugal device, and filtered through a 0.22 µm filter. Proteins were further purified via SEC on a Superdex 200 Increase 10/300 GL column (#28990944, Cytiva), concentrated, aliquoted, flash-frozen, and stored at −80°C until further use. Prior to nanoparticle assembly, endotoxin levels were confirmed to be below 50 EU/mL.

### *In vitro* nanoparticle assembly

SS-I53-50NPs were produced as previously described^37^. NPs were assembled on ice by mixing SS-I53-50A and I53-50B.4PT1 components at a final concentration of 50 µM in a 1.1:1 molar ratio and incubated for 1 h at RT with gentle rocking. Next, assembled NPs were filtered and purified by SEC using a Superose 6 Increase 10/300 GL column on an ÄKTA Pure system (Cytiva) in running buffer (1× PBS, 5% glycerol). Assembled NPs eluting in the void volume were pooled, concentrated, and flash-frozen for storage at −80°C until further use. NP concentration was determined using a previously reported formula^110^ based on absorbance at 280 nm measured with a NanoDrop Lite spectrophotometer, molecular weight, and extinction coefficient calculated using the ExPASy ProtParam tool (https://web.expasy.org/protparam/).

## SDS PAGE analysis

Immunogen purity and size were analyzed by SDS–PAGE. Briefly, 40 µL of immunogen was mixed with 10 µL ß-mercaptoethanol–supplemented Laemmli loading dye (1×) (#1610747, Bio-Rad) and heated at 95°C for 10 min. Then, 10 µL of the immunogen–dye mixture and 5 µL of PageRuler Plus Prestained Protein Ladder (#26619, ThermoFisher) were loaded into wells of a SurePAGE Bis-Tris 4–20% gel (#M00656, GenScript). Gels were run in MOPS buffer (#M00138, GenScript) for 1 h at 130 V. After electrophoresis, gels were stained with Coomassie blue for 10 min, destained, and imaged with a UV ChemiDoc MP Touch Imaging System (Bio-Rad, Waltham, MA, USA).

## Dynamic light scattering (DLS)

To measure the hydrodynamic diameter (Dh) and polydispersity of SS and NPs in solution, a Wyatt DynaPro Plate Reader III (Wyatt Technology Corp., Santa Barbara, CA, USA) was used. Measurements were performed in triplicate at 25°C in a Greiner Bio-One 96-well Sensoplate (#655892, Greiner Bio-One). Proteins diluted to 250 µg/mL were filtered and aliquoted into wells for ten acquisitions. Data were analyzed using the instrument software.

## Conventional differential scanning fluorimetry (DSF)

Protein thermal shifts were monitored using a real-time PCR instrument, QuantStudio 6/7 Flex (ThermoFisher, Waltham, MA, USA), which measures variations in fluorescence of a dye that binds to proteins as they unfold. Briefly, a MicroAmp FAST Optical 96-Well plate (#4346907, ThermoFisher) was filled with duplicates of all immunogens at 250 µg/mL and 5× SYPRO Orange Protein Gel Stain (#S6650, ThermoFisher). Next, the plate was sealed with MicroAmp optical adhesive film (#4360954, ThermoFisher). Continuous fluorescence measurements at excitation/emission wavelengths of 465/580 nm were recorded across a linear temperature range from 25°C to 95°C at a ramp rate of 1°C/min. Data were fitted using the instrument software and plotted as the first derivative of the melting curves. The peaks of the first derivative were used to determine the melting temperatures.

## Structural characterization

### Negative-stain transmission electron microscopy (NS-TEM)

Purified bare and SS-NPs were diluted to 70 µg/mL in 1× PBS, and 4 µL of the samples were applied to freshly glow-discharged carbon-coated 400-mesh copper grids (#FCF400-Cu-50, Electron Microscopy Sciences) for 1 min. The grids were blotted, dipped in 20 µL of distilled water, and blotted with Whatman no. 1 filter paper. The grids were then stained with 10 µL of 0.75% (w/v) uranyl formate, blotted to remove excess stain, and air-dried for 1 min before storage or imaging. The grids were screened and imaged using a Philips CM10 transmission electron microscope (ThermoFisher, Waltham, MA, USA) equipped with a tungsten filament and a Gatan UltraScan 894 (2k × 2k) CCD camera operating at 100 kV. Micrographs were recorded at 52,000× magnification.

### Cryo-electron microscopy (cryo-EM), processing, and 3D reconstruction

To obtain cryo-EM particle reconstructions, 4 µL of purified NP samples (SS-L2-ferritinNP, SS-L2-LuSNP, and SS-L2-I53-50NP) at 1 mg/mL were loaded onto freshly glow-discharged copper grids (1.2 µm/1.3 µm, 400 mesh, Quantifoil). The grids were blotted for 4 s at 4°C and 100% humidity, plunge-frozen using a Vitrobot Mark IV (ThermoFisher, Waltham, MA, USA), rapidly cryocooled in liquid ethane, and stored in liquid nitrogen until imaging. The grids were imaged at 1.1 Å per pixel using a Talos Arctica cryo-electron microscope (ThermoFisher, Waltham, MA, USA) operated at 200 kV. Micrographs were recorded on a post-GIF Gatan K3 camera using SerialEM version 4.1. Collection parameters, including total dose (e⁻/Å²), number of frames (e⁻/Å² per frame), defocus range, and exposure time for each NP, are listed in **Supplementary Table 2**.

The collected dataset of stacked movies was processed using the cryoSPARC v4.6.2 package^111^ as follows and detailed in **Extended Data Fig. 4–6**. Briefly, all movies were imported into cryoSPARC v4.1 and motion-corrected using the patch motion correction module. The contrast transfer function (CTF) was evaluated using patch CTF estimation. Next, after iterative rounds of manual particle picking, template-based, and automated 2D classification, particles were selected for *ab initio* 3D reconstruction (using C1 symmetry), and homogeneous refinement was performed with respective icosahedral and octahedral symmetries. All symmetries yielded cryo-EM maps with well-resolved ferritin and LuS NP cores and stable SS protruding from the core, except for I53-50NPs, in which SS was not visible. The cryo-EM maps were visualized and docked with atomic models using UCSF ChimeraX.

## Biolayer interferometry (BLI)

The antigenicity of all immunogens was assessed using a BLI detection system (Octet RH96 System, Sartorius-ForteBio) following the manufacturer’s protocol. Briefly, mAbs diluted to 40 µg/mL in 1× HBS-EP+ kinetic buffer (#BR100669, Cytiva) were added to a solid black 384-well microplate (#18-5076, Sartorius) at 100 µL per well. Immunogens were also diluted to 100 µg/mL in kinetic buffer, serially diluted twofold to a final concentration of 6.25 µg/mL, and added to the plate. Next, all biosensors were hydrated in kinetics buffer for 10 min before use. Assays were performed on the Octet RH96 system at 30°C, shaking at 1,000 rpm, as follows: hydrated Protein A biosensors (#18-5010, Sartorius) were dipped into mAbs during the loading step for 180 s, followed by a 60 s baseline step in kinetics buffer to wash off unbound mAbs. Next, immobilized mAb–biosensor tips were dipped into immunogens for 180 s during the association step and then moved back into kinetics buffer for an additional 300 s during the dissociation step. Data were acquired using Octet BLI Discovery Software v8.0 (Sartorius, Cambridge, MA, USA) and analyzed with Octet Analysis Studio Software v8.0 (Sartorius, Cambridge, MA, USA). Data were baseline-subtracted and fitted with a 1:1 binding model to calculate kinetic parameters. All experiments were independently performed in triplicate, and the values shown represent the mean.

## Serum competition BLI

Serum Ab competition was performed using an Octet RH96 instrument (Sartorius Corporation,Bohemia, NY, USA) as described previously^19,71^ to determine the proportion of Abs in the polyclonal sera. Briefly, MERS SS protein diluted in 1× HBS-EP+ kinetic buffer (#BR100669, Cytiva) at 10 μg/mL was immobilized onto equilibrated anti-Penta-HIS biosensors (Octet HIS1K, #18-5120, Sartorius) for 5 min, followed by washing in kinetic buffer to remove excess SS proteins. Next, SS-immobilized probes were transferred into serially diluted sera from vaccinated mice or kinetic buffer for 5 min and then dipped into 50 µg/mL competing mAbs for 3 min to assess competitive serum Ab inhibition. All sera were diluted fourfold, eight times in kinetic buffer, starting at 1:10. The raw data for Ab competitive binding signals were analyzed and fitted with a 1:1 binding model in Octet software v8.0, and results were plotted using GraphPad Prism v10.3.0. A representative plot is shown in **Supplementary Fig. 1b**. Ro represents the response of the maximum noncompeting binding curve, and Rc represents the response of the serum-competing binding curve. The formula ([Ro − Rc] × 100 / Ro) was used to calculate the percentage of inhibition or competition. All experiments were independently performed in duplicate, and the data shown represent the mean.

## Immunogen–adjuvant preparation

Immunogen–adjuvant formulations were prepared as previously described^5^. Briefly, purified immunogens low endotoxin concentrations (<50 EU/mL) were diluted to provide 10 μg per 50 μL dose upon addition of adjuvant. An oil-in-water emulsion adjuvant from the Sigma Adjuvant System (SAS; #S6322, Sigma) was used. Before formulation, SAS adjuvant was warmed to 42°C and mixed with sterile 1× PBS (pH 7.4) following the manufacturer’s recommendations. Immunogens at the appropriate dose or control 1× PBS (pH 7.4) were mixed with the prepared SAS adjuvant at a 1:1 volume ratio, and the mixtures were vortexed for 1 min. Prior to immunization, the immunogen–adjuvant mixtures were thoroughly mixed on a shaker for 20 min at 4°C.

## Endotoxin assessment and removal

Endotoxin levels were assessed and removed as previously described^37^. For I53-50B.4PT1 produced in bacteria, endotoxin was first removed during Ni-NTA affinity purification by prolonged washing with detergent-containing buffer (1× PBS, 0.75% CHAPS). Purified I53-50B.4PT1 (before use in I53-50NP assembly) and all immunogens were tested for endotoxin using the Kinetic-QCL™ Kinetic Chromogenic Limulus Amebocyte Lysate (LAL) kit (#50-650U, Lonza Bioscience) following the manufacturer’s recommendations. The measured endotoxin concentrations were routinely negative or below <50 EU/mg, a threshold suitable for immunization.

## Mouse immunizations

Immunizations were performed using standard schedules and procedures as previously described^21^. Briefly, BALB/cJ mice, as described above, received two IM injections at weeks 0 and 3 with 50 µL per hind leg. Two weeks post-injection, mice were euthanized, and blood was collected via terminal cardiac puncture. Blood samples were allowed to clot for 30 min; sera were collected and stored at −80°C or heat-inactivated at 56°C for 1 h to evaluate immune responses via subsequent serological assays.

## Anti-sera enzyme-linked immunosorbent assay (ELISA)

ELISA was used to determine serum binding Abs to different S-2P proteins as previously described^104^. Briefly, 96-well EIA/RIA clear flat-bottom microplates (#33619, Corning) were coated with 1 µg/mL of protein and incubated at 4°C for 16 h. After standard washes in PBS-T buffer (1× PBS + 0.05% Tween 20) and blocking with blocking buffer (PBS-T + 5% nonfat Difco skim milk, #232100, Midland Scientific) for 1 h at RT, plates were incubated with serial dilutions of heat-inactivated mouse sera (starting at 1:100) for 1 h at RT. Where applicable, 50 μg/ml of foldon protein was added for 1 h at RT to block foldon-specific Ab binding. Following washes, a 1:4000 dilution of goat anti-mouse IgG (H+L) horseradish peroxidase conjugate (#G-21040, ThermoFisher) in blocking buffer was added and incubated for 1 h at RT. Next, plates were rewashed and developed with 100 μL of 1-Step 3,5,3′,5′-tetramethylbenzidine ELISA peroxidase substrate solution (#N301, ThermoFisher) to detect Ab responses. Plates were quenched with 100 µL of 1 N sulfuric acid (#SA212-1, ThermoFisher) and read at 450 nm and 650 nm using a SpectraMax iD5 Multi-Mode Microplate Reader (Molecular Devices, San Jose, CA, USA). Endpoint titers were defined as the highest dilution showing absorbance (A450) fourfold above the background (secondary Ab only).

## Pseudotyped lentivirus production

Pseudotyped lentiviral reporter viruses were produced as previously described^5,104^. Briefly, pseudoviruses bearing codon-optimized S glycoproteins of MERS-CoV, SARS-CoV-2, and WIV1-CoV, along with a firefly luciferase (Luc) reporter gene, were produced by co-transfecting HEK293T/17 (ATCC CRL-11268) cells with 17.5 µg of pCMVΔR8.2 (lentiviral backbone), 17.5 µg of pHR’CMVLuc (luciferase reporter), and 1 µg of spike (S gene) plasmids seeded in 150 mm tissue culture dishes using FuGENE 6 transfection reagent (#E2692, Promega) in Opti-MEM Reduced Serum Medium (Gibco #31985-070). For SARS-CoV-2 and WIV1-CoV pseudoviruses, 0.31 µg of pCMV-TMPRSS2 (human transmembrane protease serine 2) plasmid was also co-transfected. Following overnight incubation, dishes were replenished with fresh complete medium. Next, pseudovirus supernatants were harvested 48 h later, filtered through a 0.45 µm low-protein-binding Steriflip PVDF filter (#SE1M003M00, Sigma), and frozen at −80°C. Each pseudotyped viral stock was titrated before use in neutralization assays by infecting target cells for 72 h, and luciferase activity was determined using the Luciferase Assay System (#E1501, Promega). Pseudovirus titers are expressed as relative luminescence units (RLU).

## MERS-CoV pseudotyped virus neutralization assay

MERS-CoV pseudotyped virus neutralizing activity was evaluated as previously described^5,29^. Briefly, Huh7.5 cells were seeded in 96-well black and white Isoplates (#6005068, Revvity) at 10,000 cells/well and incubated at 37°C with 5% CO₂ overnight. The next day, sera samples were serially diluted (starting at 1:40, 4-fold, ×8) in 90 μL DMEM media in U-bottom MicroWell plates (#249946, ThermoFisher), and 90 µL of pseudovirus expressing MERS-CoV England1 spike (at 100,000 RLU) was added. The sera–pseudovirus mixtures were incubated at 37°C for 45 min, and 50 μL was added to the seeded cells in triplicate. The cells were then incubated at 37°C with 5% CO_2_ for 2 h to allow infection. Plates were then topped up with 100 µL of DMEM supplemented with 10% FBS and 1% penicillin–streptomycin and incubated under the same conditions for 72 h. Pseudovirus infectivity or neutralizing activity was determined by measuring luciferase activity using the Luciferase Assay System. Cells were lysed, and 50 µL of luciferase substrate was added. Luminescence was read at 570 nm (RLU) using a SpectraMax iD5 Multi-Mode Microplate Reader. Percent neutralization was normalized, considering uninfected cells as 100% neutralization and cells transduced with pseudovirus alone as 0% neutralization.

Triplicate averages were normalized, and sigmoidal curves were generated. NAb titers at 50% (ID50) were determined by applying a [log(agonist) vs. normalized-response (variable slope)] nonlinear regression model to normalized values in GraphPad Prism v10.3.

## SARS-CoV-2 and WIV1-CoV pseudotyped virus neutralization assay

SARS-CoV-2 and WIV1-CoV pseudotyped virus neutralizing activity was determined similarly to the MERS-CoV pseudovirus neutralization assay, with minor modifications as previously described^21,104^. Briefly, MF_293T-hACE2^104^ (provided by M. Farzan, Scripps Research Institute) were seeded in 96-well black and white Isoplates at 5,000 cells/well and incubated at 37°C with 5% CO_2_. Sera samples, diluted similarly as for the MERS-CoV pseudovirus neutralization assay, were mixed with pseudoviruses expressing SARS-CoV-2 JN.1 or WIV1-CoV spikes and incubated at 37°C with 5% CO_2_ for 45 min. The sera–pseudovirus mixtures were added to the seeded cells in triplicate and incubated for 2 h to allow infection. Plates were then topped up with 100 µL of DMEM supplemented with 10% FBS, 1% penicillin–streptomycin, and 3 µg/mL puromycin, and incubated under the same conditions for 72 h. Luciferase readouts were determined using the same methods as described above for the MERS-CoV pseudovirus neutralization assay.

## Antibody isotyping and Fc receptor binding characterization

Mouse sera were analyzed using a custom-developed Luminex-based multiplex immunoassay to assess antigen-specific Ab features, following previously established methodologies^73^. A panel of antigens—including prefusion-stabilized S proteins from MERS-CoV, SARS-CoV, SARS-CoV-2, WIV1-CoV, HCoV-HKU1, HCoV-OC43, stabilized MERS-CoV and SARS-CoV-2 SS and EBOV GP (used as a negative control antigen)—were covalently conjugated to distinct MagPlex bead regions using EDC (#A35391,ThermoFisher) and Sulfo-NHS (#A39269, ThermoFisher). Once conjugated, the antigen-coupled beads were blocked and pooled for multiplexing. To determine optimal detection ranges, preliminary titrations were conducted to identify the appropriate serum dilution for each Ig isotype and Fc receptor target. Samples were then diluted and incubated with the pooled bead mixture in 384-well plates (in duplicate) for 2 h at RT in Assay Buffer (0.1% BSA and 0.02% Tween 20 in 1× PBS, pH 7.4) with orbital shaking at 750 rpm to facilitate immune complex formation. Plates were subsequently washed three times with Assay Buffer using a magnetic Tecan Hydrospeed Plate Washer.

The wash cycle included the following steps: aspiration, soak (30 s), aspiration, dispense (60 µL at 100 µL/s), soak and shake (medium intensity, 10 s), soak (40 s), and final aspiration (1 s at 4.5 mm), performed for three cycles as specified. To detect specific immunoglobulin isotypes, PE-conjugated goat anti-mouse antibodies (IgG1, IgG2a, IgG3, IgM, and IgA; Southern Biotech) were diluted in Assay Buffer and added to the plates, followed by a 1 h incubation at RT under the same shaking conditions. For Fc receptor profiling, recombinant mouse Fcγ receptors (FcγRIIb, FcγRIII, and FcγRIV; produced at Duke University, Durham, NC, USA) were biotinylated using the BirA500 kit (#NC2047451, Avidity) and pre-incubated with Streptavidin-PE before application to the wells. Plates were incubated for an additional 1 h and then washed again with Assay Buffer for three washes. Finally, beads/immune complexes were resuspended in 1× Luminex Sheath Reagent for acquisition on the Luminex xMAP INTELLIFLEX DR-SE platform. MFI values were calculated using the system software, and each data point represents the mean of technical duplicates. All reported values were quality controlled as being above the background (no serum added). Naïve responses were shown as reference controls and were also background subtracted. Technical replicates whose standard deviation was greater than 50% of the mean were not reported.

## Functional characterization of Fc-mediated antibody effector mechanisms

### Antibody-dependent cellular and neutrophil phagocytosis (ADCP and ADNP)

Phagocytosis assays were conducted using a flow cytometry–based method. Antigens identical to those used in the Ab binding assay were biotinylated with EZ-Link Sulfo-NHS-LC-LC-Biotin (#21338, ThermoFisher), purified using Zeba desalting columns, and conjugated to yellow/green fluorescent NeutrAvidin microspheres (#F8776, ThermoFisher) using 0.1% BSA in 1× PBS, pH = 7.4. Conjugated beads were transferred to 96-well round-bottom plates and incubated with 1:100 diluted mouse sera at 37°C for 2 h to allow immune complexes to form. Plates were then washed with PBS, centrifuged at 200 × g for 10 min, and the supernatant was discarded. For ADCP, 25,000 monocytes per well were added; for ADNP, 50,000 neutrophils were added. Cells were incubated for 2 h at 37°C. Next, plates were centrifuged (500 × g, 5 min), and unbound material was removed. For cell staining, CD14 PacBlue Ab (#305112, BioLegend) for ADCP or CD66b V450 Ab (#305112, BioLegend) for ADNP were added at 1:100 dilution and incubated for 10 min at RT. After washing (centrifugation at 500 × g for 5 min and supernatant discarded), cells were fixed in 4% paraformaldehyde (PFA) for 10 min, washed again, and resuspended in PBS for analysis on the iQue 3 HTS Cytometer (Sartorius, Cambridge, MA, USA). Gated monocyte and neutrophil populations were evaluated for fluorescent bead uptake to calculate phagocytic scores.

Monocytes for ADCP were obtained from human peripheral blood leukopaks (#70500.2, Stemcell) via red blood cell (RBC) depletion using an EasySep RBC Depletion Kit (#18170, Stemcell). EDTA (#15575-038, ThermoFisher) was added to preserve cell viability, followed by incubation with depletion reagent and separation using an EasySep magnet. This step was repeated to ensure removal of residual RBCs. A gating strategy for CD14+ monocyte ADCP is shown in **Supplementary Fig. 2a**.

Neutrophils for ADNP were isolated from fresh human whole blood (#70500.2, Stemcell) using the EasySep Direct Human Neutrophil Isolation Kit (#19666, Stemcell). Blood was treated with an isolation cocktail and RapidSpheres, diluted with EasySep buffer (#20144, Stemcell), and subjected to magnetic separation twice to maximize purity, leaving behind all RBCs. A gating strategy for CD66+ neutrophil ADNP is shown in **Supplementary Fig. 2b**. The use of human cells to validate murine opsonophagocytosis responses has been previously validated and reported^82^.

### Antibody-dependent complement deposition (ADCD)

ADCD assays were performed with the same antigens (used in Ab binding) conjugated to MagPlex beads as previously described^112^. The antigen–bead complexes were mixed and dispensed into 384-well plates containing 1:20 diluted serum samples (in duplicate) and incubated for 2 h at RT in Assay Buffer with shaking at 800 rpm. Following incubation, plates were washed with 1% BSA in PBS using a magnetic Tecan Hydrospeed washer. Lyophilized guinea pig complement (#CL4051, Cedarlane), reconstituted in gelatin veronal buffer with calcium and magnesium (#G6514, Sigma), was added and incubated for 20 min at 37°C with shaking at 800 rpm. Plates were rewashed with 0.1% BSA in PBS. C3 complement deposition was detected by adding FITC-labeled anti–guinea pig C3 Ab (#0855385, MP Biomedicals), incubating for 30 min at RT, and performing a final wash. Beads were resuspended in PBS and analyzed on the iQue 3 HTS platform. MFI values were calculated and averaged across replicates. A gating strategy for ADCD is shown in **Supplementary Fig. 2d**.

### Antibody-dependent natural killer cell activation (ADNKA)

ADNKA was quantified via the surface expression of CD107a (as a marker for degranulation) and MIP-1β (as a marker for NK cell activation). ELISA plates were coated with 3 µg/mL of the same antigens used in binding assays and incubated at 37°C for 2 h. Plates were washed with 1× PBS, blocked with 5% BSA in 1× PBS for 1 h at 37°C, and washed again. Mouse serum diluted 1:25 in PBS was added and incubated overnight at 4°C. The next day, plates were washed three times with 1× PBS. NK cells (prepared as described below) were added to the antigen-coated plates and incubated for 5 h at 37°C with 5% CO₂. After incubation, cells were transferred to V-bottom 96-well plates preloaded with a surface stain cocktail containing CD56 PE-Cy7 (#557747, BD Biosciences), CD16 APC-Cy7 (#557758, BD Biosciences), and CD3 PacBlue (#558117, BD Biosciences).

Plates were stained for 15 min in the dark at RT, washed twice (centrifugation at 500 × g, 5 min), and fixed with PermA (#GAS001S100, ThermoFisher) for 15 min under foil. Following fixation, intracellular staining with anti–MIP-1β PE (#550078, BD Biosciences) in PermB buffer (#GAS002S100, ThermoFisher) was performed for 15 min. Cells were resuspended in PBS and analyzed on the iQue 3 HTS platform. NK cells were gated as CD56⁺/CD16⁺/CD3⁻. The percentage of NK cells expressing CD107a, MIP-1β, and IFNγ was measured as a readout of NK activation. Assays were performed in duplicate using NK cells from at least two healthy donors to ensure reproducibility. A gating strategy for ADNKA is shown in **Supplementary Fig. 2c**.

Natural killer cells were isolated from fresh leukopaks (#70500.2, Stemcell) using the EasySep Human NK Cell Isolation Kit (#17955, Stemcell). Following centrifugation, the supernatant was extracted, and cells were resuspended to 5 × 10⁷ cells/mL. The isolation cocktail was added, followed by a 5 min incubation at RT. Magnetic RapidSpheres were then added, and the sample was diluted to 50 mL with buffer, mixed, and placed in the magnet for 5 min. The supernatant containing enriched NK cells was transferred to a new tube and re-exposed to the magnet to enhance purity. NK cells were cultured in R10 medium—RPMI 1640 (#11875093, ThermoFisher), 10% FBS, 1% penicillin/streptomycin, and 4 mM L-glutamine—with IL-15 overnight at 37°C with 5% CO₂ at a density of 1.5 × 10⁶ cells/mL. Prior to use, cells were adjusted to 2.5 × 10⁵ cells/mL and incubated with a cocktail of anti-CD107a PE-Cy5 (#555802, BD Biosciences), Brefeldin A (#B7651, Sigma), and GolgiStop (#554724, BD Biosciences). The use of human cells to validate murine NK responses has been previously validated and reported^82^.

## Quantification and statistical analysis

Details of statistical experiments, including all quantifications, statistical analyses, tests, and animal numbers (N), are included in the figure legends. Experiments were conducted in triplicate using sera from ten BALB/cJ animals for anti-sera ELISAs and neutralization assays. Prior to statistical analysis, all values were log₁₀-transformed after reciprocal endpoint and reciprocal ID_50_ titers were calculated. Geometric mean titers were determined. Assay limits of detection (LOD) are indicated by dotted lines. Two-way ANOVA and Tukey’s multiple comparisons were used to evaluate the datasets. To ascertain whether our data were normally distributed—a prerequisite for Tukey’s multiple comparison tests—we compared actual and projected binding and neutralization titers using quantile–quantile probability plots. All data followed a normal distribution. Statistical analyses were performed using GraphPad Prism v10.3.0. Significance is indicated with asterisks defined as *P < 0.05, **P < 0.01, ***P < 0.001, ****P < 0.0001 and non-significant groups are not shown. Systems serology quantifications were performed using the mean of technical replicates (binding assays) or biological donors (Fc-effector assays) and, where indicated, subtracting baseline values (no serum) for each feature. All data were quality controlled to ensure that standard deviations of the replicates did not exceed 50% of the mean; if replicates exceeded that threshold, the data were not reported. For all binding assays, serum samples whose binding features did not exceed baseline values (no serum) + 3 standard deviations were not considered reportable. For all Fc-effector assays, baseline values were directly subtracted from the reported value. Statistical analyses for systems serology–generated data were performed using ANOVA with Tukey’s post hoc test. Primary comparisons were each vaccinated group versus the naïve cohort. Secondary comparisons were between nanoparticle groups.

## Data and code availability

Source data supporting the findings of this study are available within this paper and its supplementary files. Source data has also been deposited to Figshare and can be accessed here: 10.6084/m9.figshare.30454586. Additional source data inquiries should be directed to the corresponding author, Kizzmekia Corbett-Helaire. Cryo-EM reconstructions for SS-L2-FerritinNP, SS-L2-LuSNP and SS-L2-I53-50NP have been deposited to Electron Microscopy Data Bank under accession codes EMD-73668, EMD-73671, and EMD-73675, respectively. No codes were used in data acquisition or analysis for this paper. Systems serology data were analyzed through a customized R-based pipeline, and all data processing and code were performed using R version 6.0.

## Material and resource availability

All requests for reagents and resources should be directed to the corresponding author, Kizzmekia Corbett-Helaire and will be made available after completion of a Material Transfer Agreement (MTA). If the material was obtained under use restriction, the inquiry will be forwarded to appropriate party.

## Acknowledgements

We thank Fidan Baycora for administrative support and Emily Hobbs for grant support. We express our sincere gratitude to the Harvard Cryo-Electron Microscopy Center for Structural Biology, Harvard Medical School’s Molecular Electron Microscopy Suite, and the Center for Macromolecular Interactions for providing services and access to equipment and facilities. We also thank all members of the Harvard Center for Comparative Medicine mouse facility at the Harvard T.H. Chan School of Public Health for animal husbandry assistance. We thank Nicole Doria-Rose and NIH, NIAID, VRC’s Humoral Immunology Core for pseudovirus neutralization assay standard operation procedure training. This work was supported in part by the Howard Hughes Medical Institute Freeman Hrabowski Scholars grant (to KSC), the Blavatnik Biomedical Accelerator at Harvard University (to KSC), the Hans Sigrist Foundation Prize (to KSC), in-kind gifts of lab equipment, consumables, and supplies from Corning, Inc. (to KSC), and the Melvin J. and Geraldine L. Glimcher Assistant Professorship and start-up funds from the Harvard T.H. Chan School of Public Health (to KSC). Support for systems serology assays and analysis was provided by NIAID (P01AI65072 and U19AI135995 to RPM). AL’s MD-PhD training is supported by NIH National Institute of General Medical Sciences (T32GM144273).

## Author contributions

CKOD and KSC conceptualized the study. CKOD designed linker sequences, engineered and produced NP immunogens. CKOD and SPM performed protein affinity purification. CKOD performed all SEC purification and analysis. CKOD performed NS-TEM, DLS, DSF, cryo-EM, and BLI antigenicity experiments and analysis. CKOD, SPM, and KSC designed animal immunization experiments; animal experiments were completed by CKOD and SPM. SPM, AT, and OA performed ELISAs. CKOD and SPM performed lentiviral pseudovirus production and neutralization assays. SPM, AT, and VF performed pseudovirus neutralization data processing. CKOD and AL optimized the serum competition BLI assay. CKOD, TK, and LRE performed the serum competition BLI assay. LRM performed system serology analysis under the supervision of RPM. CKOD and AED performed computational and sequence analysis. KSC supervised and administered the work. CKOD outlined and wrote the original manuscript. CKOD, SPM, and KSC revised and polished the manuscript with input from all coauthors.

