## Extended Data for "Unconventional linkers facilitate potent stabilized coronavirus stem antibody responses following nanoparticle vaccination"

### Extended Data Figures

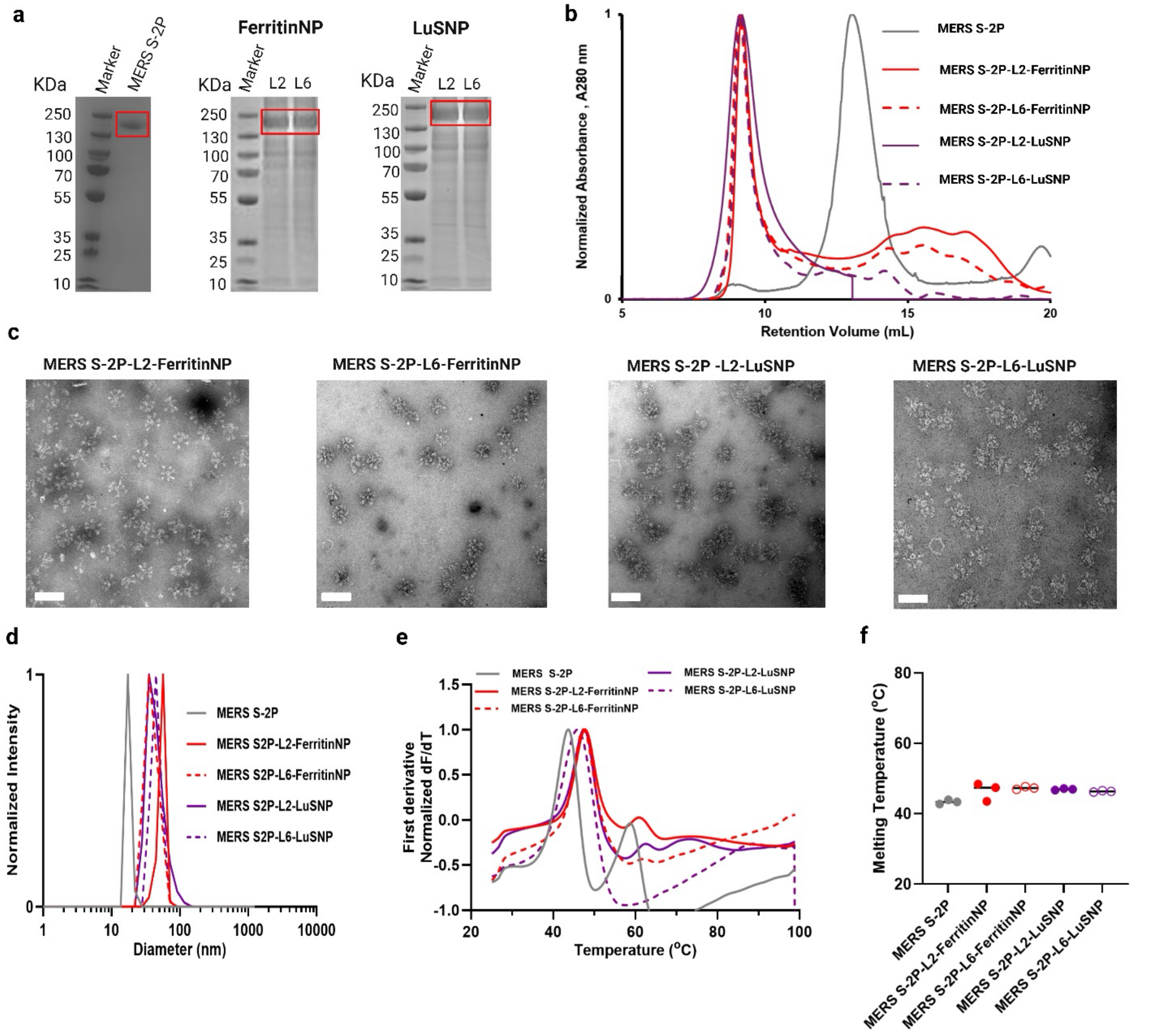


**Extended Data Fig. 1: Design, production, and characterization of MERS-CoV S-2P nanoparticles (NPs), related to Fig. 1.** (**a**) SDS–PAGE gel of size-exclusion chromatography (SEC)-purified MERS-CoV S-2P protein and affinity-purified NP immunogens. Molecular weight standards are indicated in kDa. Red boxes highlight bands corresponding to S-2P soluble protein or NPs. (**b**) Size-exclusion chromatograms of MERS S-2P soluble protein and NPs on a Superose 6 Increase 10/300 column. (**c**) Negative-stain electron micrographs (NS–EM) of purified NPs. Scale bar, 50 nm. (**d-f**) MERS soluble S-2P and NPs characterization by dynamic light scattering (DLS) (d) where hydrodynamic diameter (HD) and polydispersity index (PDI) are indicated, conventional differential scanning fluorimetry (DSF) to measure thermal melting profiles (e), and differential scanning fluorimetry to measure thermal melting temperatures (f). **(f)** Each dot represents one of three technical replicates, and bars represent means. **(b, d-f)** Colors represent MERS-CoV soluble S-2P (grey), L2-ferritinNP (red, solid line or closed circles), L6-ferritinNP (red dashed line or open circles), L2-LuSNP (purple, solid line or closed circles), L6-LuSNP (purple, dashed line or open circles).

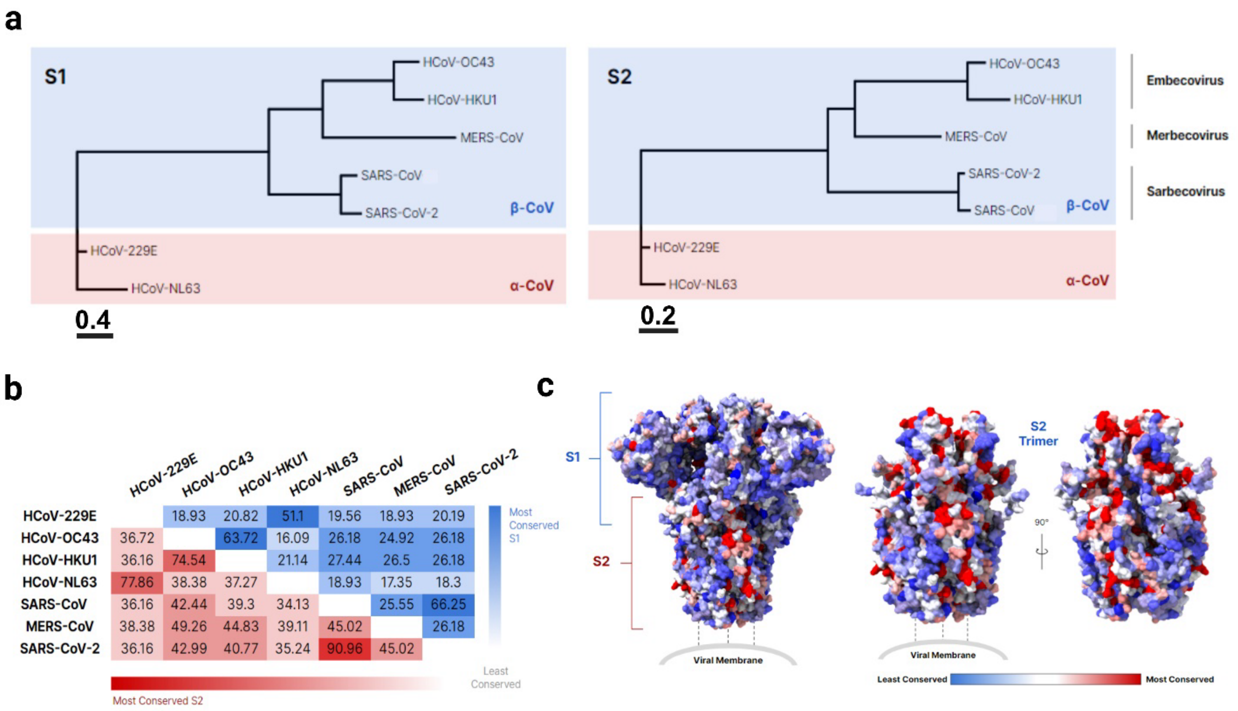


**Extended Data Fig. 2**: **Conservation and variability of human-infecting CoV S proteins, related to Figs. 1 and 2.** (**a**) Phylogenetic trees of seven human-infecting CoVs (HCoVs: HCoV-229E, -OC43, -HKU1, and -NL63 | epidemic CoVs: MERS-CoV, SARS-CoV, and SARS-CoV-2) and based on S1 (left) and S2 (right) domain sequences. Braces indicate CoV subgenera. The scale bar indicates evolutionary distance between CoVs. (**b**) Sequence identity of aforemenentioned CoVs based on S1 and S2 domain sequences. S1 conservation (top, blue). S2 conservation (bottom, red). Consensus identity is shown on a scale from red or blue (most conserved) to white (least conserved). Numbers indicate percentage identity between different HCoV S1 and S2 domains. (**c**) Structural sequence conservation from aformentioned CoVs overlaid on MERS-CoV S (PDB ID: 5W9J). Sequence alignment was generated using SARS-CoV-2 (GenBank accession QHR63290.2), SARS-CoV (GenBank accession AAP41037.1), MERS-CoV (GenBank accession ALK80242.1), HCoV-HKU1 (GenBank accession ADN03339.1), HCoV-OC43 (GenBank accession QEG03814.1), HCoV-229E (GenBank accession ABB90529.1), and HCoV-NL63 (GenBank accession QED88026.1). Sequence conservation was calculated using the ConSurf database. Blue indicates conservation. Red indicates variability.

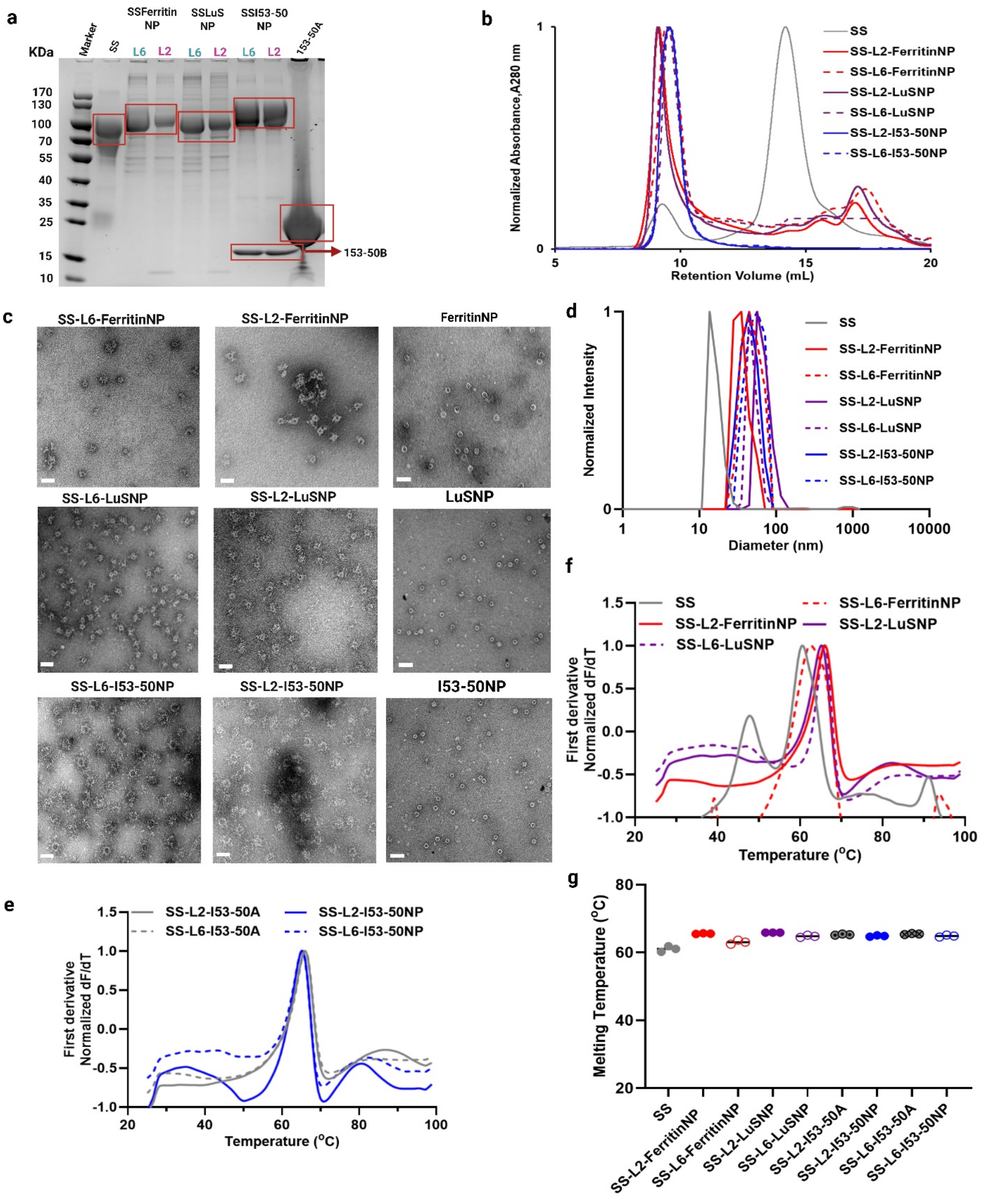


**Extended Data Fig. 3: Production and characterization of SS–NPs, related to Fig. 2.** (**a**) SDS–PAGE gel of size-exclusion chromatography (SEC)-purified MERS-CoV SS protein and affinity-purified NP immunogens. Molecular weight standards are indicated in kDa. Red boxes highlight bands corresponding to SS soluble protein or NPs. (**b**) Size-exclusion chromatograms of MERS SS soluble protein and NPs on a Superose 6 Increase 10/300 column (various colors as indicated). (**c**) NS–EM of purified NPs. Scale bar, 50 nm. (**d-g**) MERS-CoV soluble SS and NPs characterization by dynamic light scattering (DLS) (d) where hydrodynamic diameter (HD) and polydispersity index (PDI) are indicated and DSF to measure thermal melting temperatures of SS-I53-50NPs (e) and SS-ferritinNPs and SS-LuSNPs (f). **(g)** Thermal melting temperatures of immunogens measured by DSF. Each dot represents one of three technical replicates, and bars represent means. **(b, d-g)** Colors represent MERS-CoV soluble SS (grey), L2-ferritinNP (red, solid line or closed circles), L6-ferritinNP (red, dashed line or open circles), L2-LuSNP (purple, solid line or closed circles), L6-LuSNP (purple, dashed line or open circles), SS-L2-I53-50A (grey, closed dotted circles), SS-L6-I53-50A (grey, closed dotted circles), SS-L2-I53-50NP (blue, closed circles) and SS-L6-I53-50NP (blue, open circles).

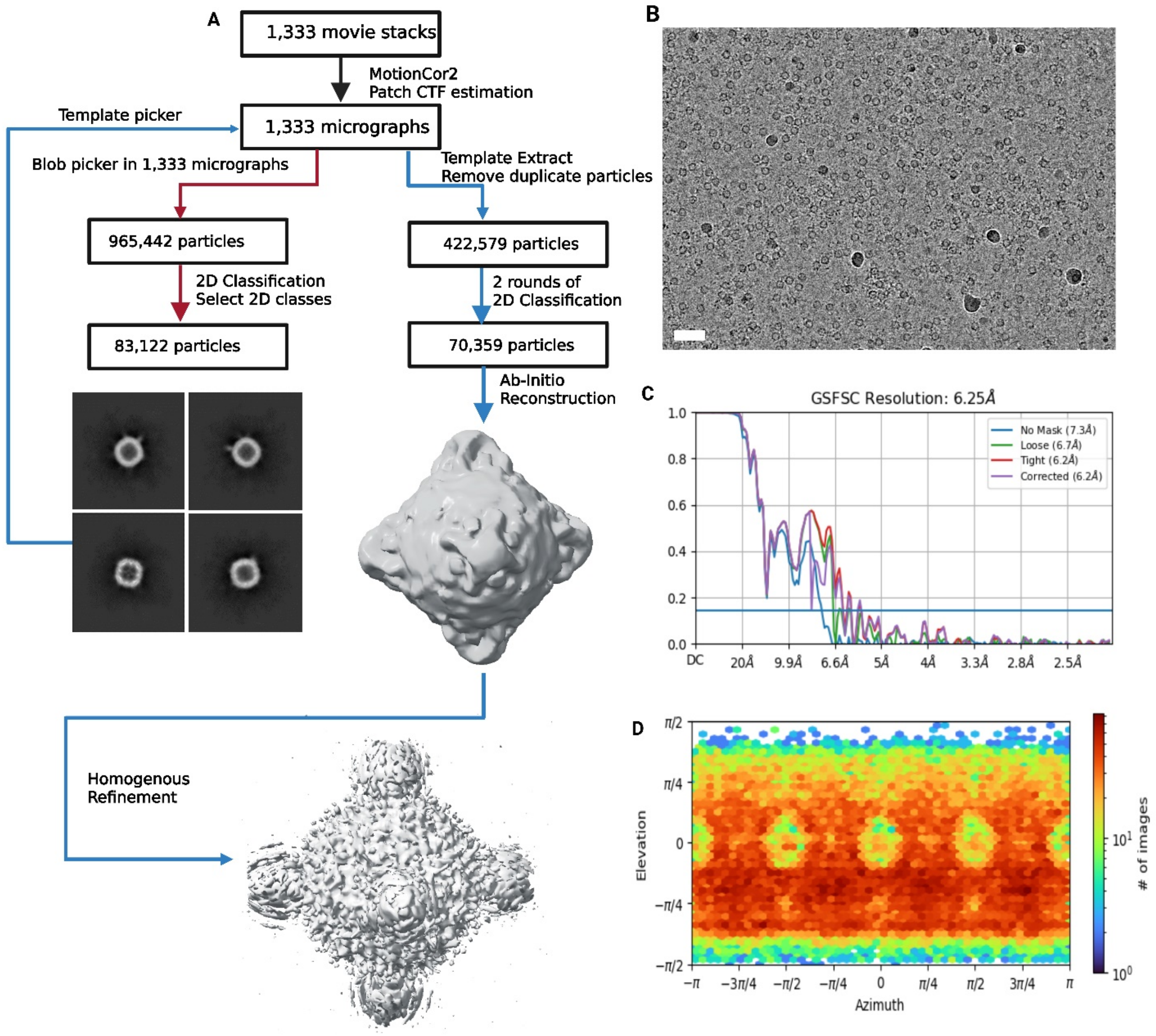


**Extended Data Fig. 4: Cryo-EM structure determination of SS-L2-ferritinNP, related to Fig. 2f.** (**a**) Cryo-EM data processing and reconstruction workflow for all relevant and standard steps. 3D reconstruction was done using octahedral symmetry. Final resolution was 6.25 Å. (**b**) Representative electron micrograph of SS-L2-ferritinNP embedded in vitreous ice. Scale bar, 20 nm. (**c**) Gold-standard Fourier shell correlation (FSC) curve for the density map. The 0.143 cut-off value is indicated by a horizontal blue bar. (**d**) 2D Euler angular distribution plots used in reconstructions.

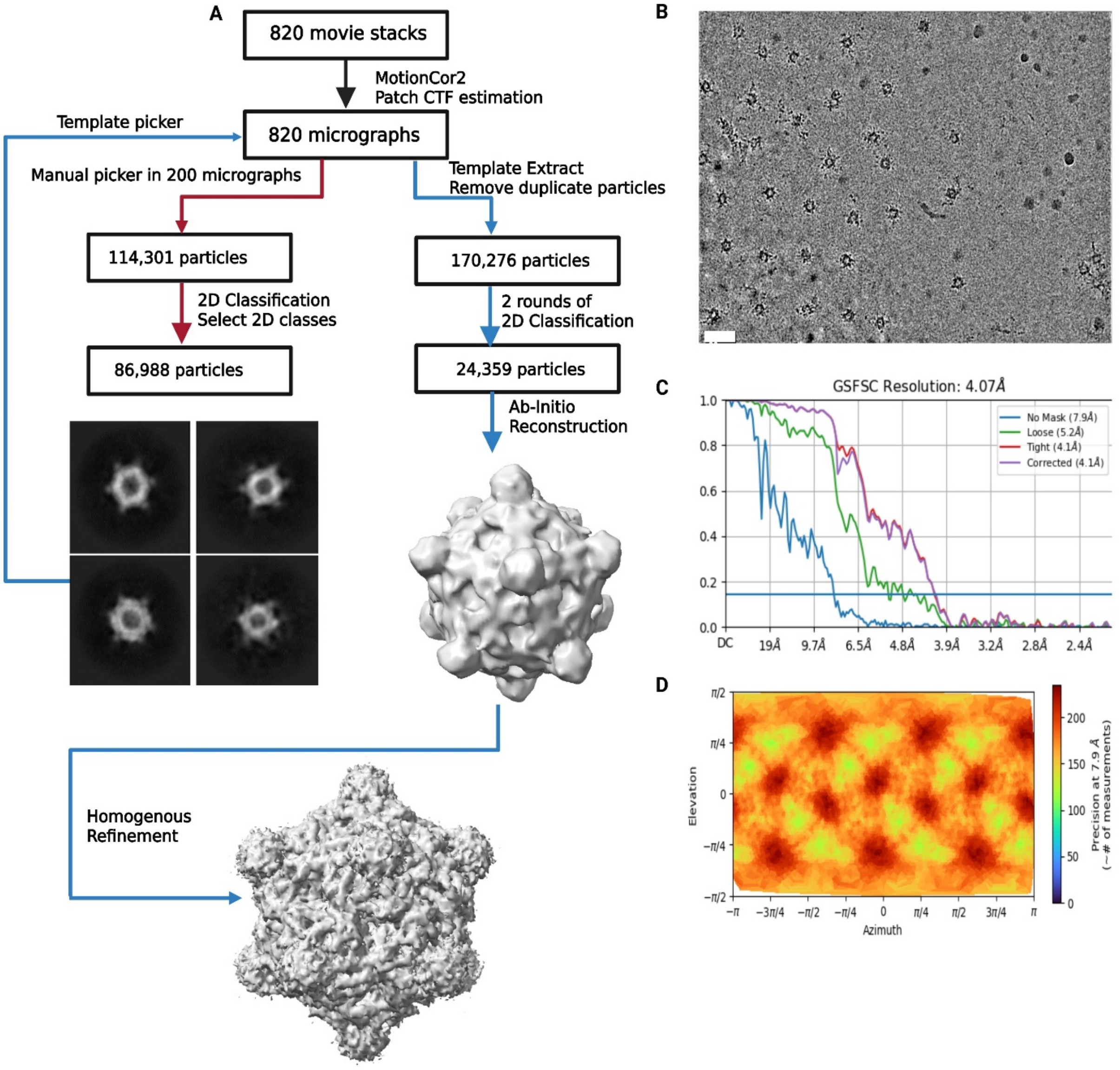


**Extended Data Fig. 5: Cryo-EM structure determination of SS-L2-LuSNP, related to Fig. 2f.** (**a**) Cryo-EM data processing and reconstruction workflow for all relevant and standard steps. 3D reconstruction was done using icosahedral symmetry. Final resolution was 4.07 Å. (**b**) Representative electron micrograph of SS-L2-LuSNP embedded in vitreous ice. Scale bar, 20 nm. (**c**) Gold-standard FSC curve for the density map. The 0.143 cut-off value is indicated by a horizontal blue bar. (**d**) 2D Euler angular distribution plots used in reconstructions.

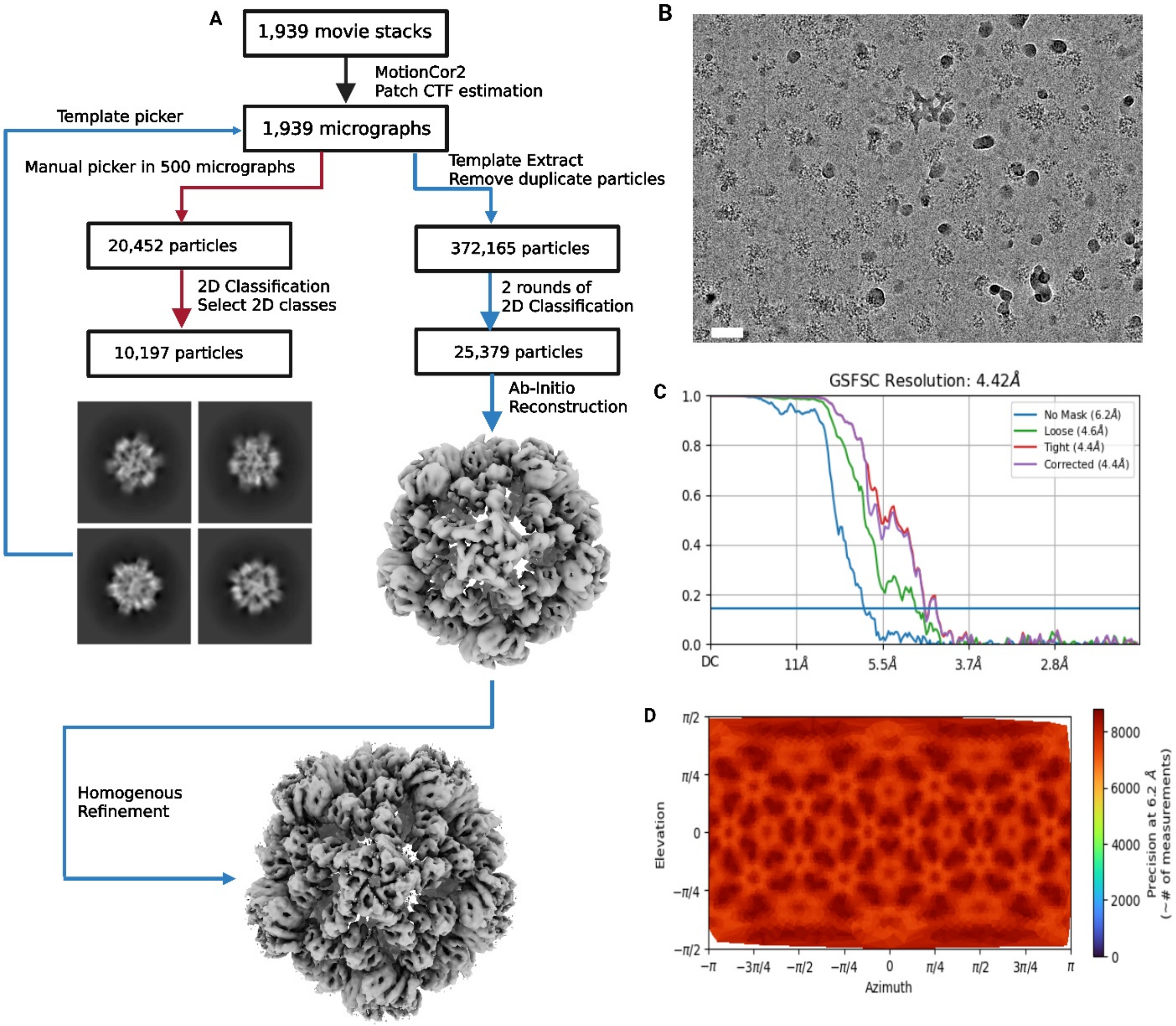


**Extended Data Fig. 6: Cryo-EM structure determination of SS-L2-I53-50NP, related to Fig. 2f.** (**a**) Cryo-EM data processing and reconstruction workflow for all relevant and standard steps. 3D reconstruction was done using icosahedral symmetry. Final resolution was 4.42 Å. (**b**) Representative electron micrograph of SS-L2-I53-50NP embedded in vitreous ice. Scale bar, 20 nm. (**c**) Gold-standard FSC curve for the density map. The 0.143 cut-off value is indicated by a horizontal blue bar. (**d**) 2D Euler angular distribution plots used in reconstructions.

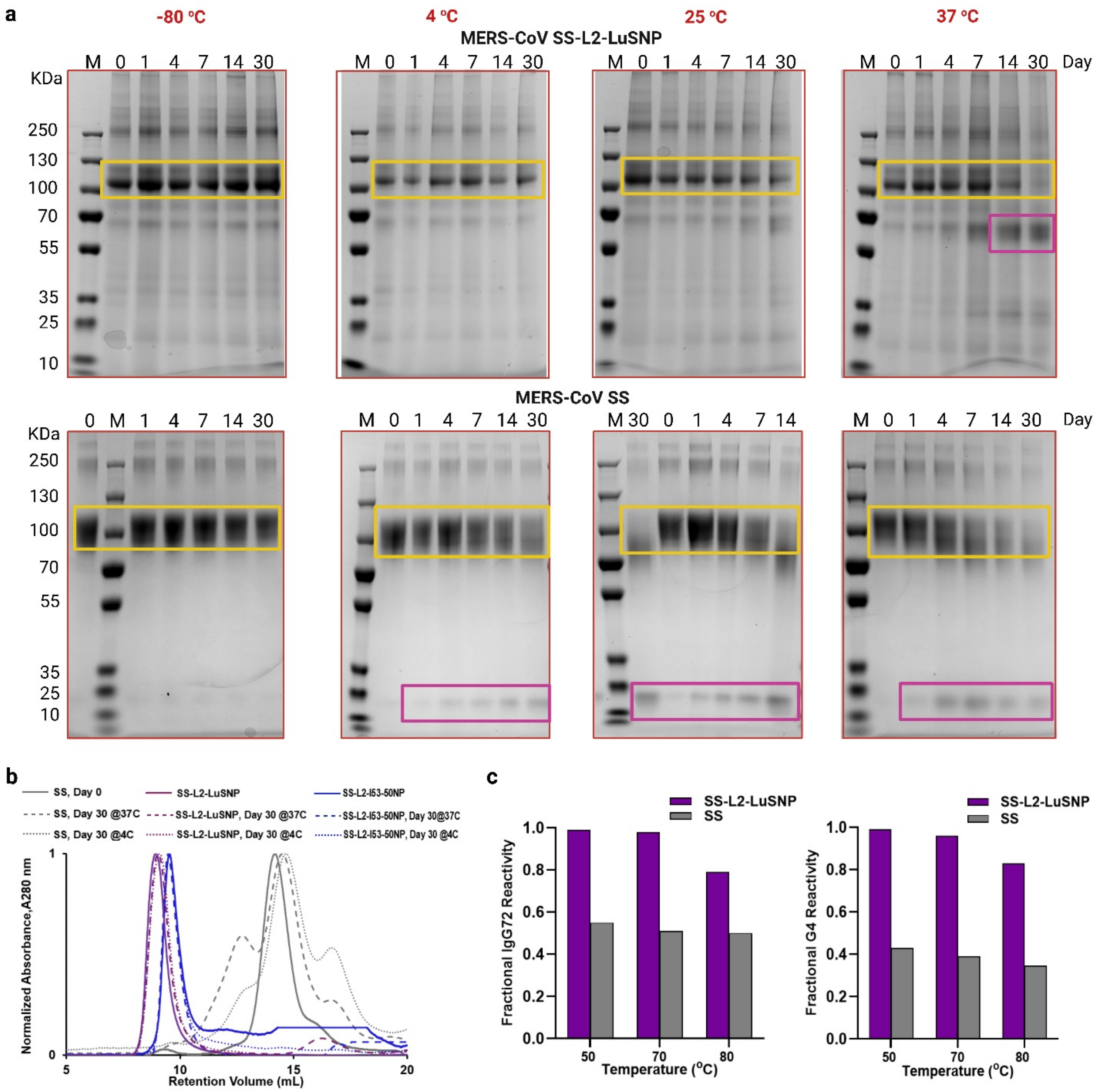


**Extended Data Fig. 7: Storage and physical stability characterization of immunogens related to Fig. 2.** (**a**) SDS–PAGE of MERS-CoV SS-L2-LuSNPs (top) and soluble SS (bottom) monitored at −80°C, 4°C, 25°C, and 37°C over a 30-day storage period. The left lane indicates molecular weight markers (M) (kDa), and subsequent lanes represent samples collected on days 0, 1, 4, 7, 14, and 30. Yellow and purple boxes highlight bands corresponding to intact and degraded proteins or NPs. (**b**) Normalized SEC profiles of MERS-CoV SS, SS-L2-LuSNP, and SS–L2–I53-50NP immunogens obtained on days 0 and 30 at 4°C and 37°C on a Superose 6 Increase 10/300 GL column. (**c**) Physical stability and retention of mAb (IgG72, left | G4, left) binding after thermal stress, measured by (BLI). Data represents three independent experiments. y-axis shows the amplitude of BLI signal obtained from immunogens incubated at the indicated temperatures for 1 h relative to the signal after incubation at 20°C.
