## Supplementary File for "Unconventional linkers facilitate potent stabilized coronavirus stem antibody responses following nanoparticle vaccination"

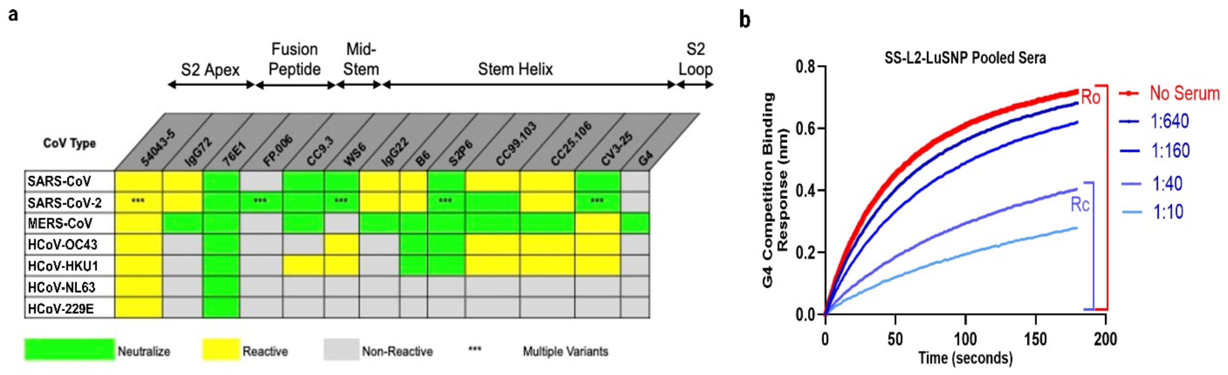


**Supplementary Fig. 1: Dissecting epitope specificity of polyclonal antibody (Ab) responses by biolayer interferometry (BLI) competition assay, related to Fig. 4** (**a**) Chart of cross-reactivity across all seven human-infecting coronaviruses (CoVs). Yellow depicts reactivity, gray indicates no reactivity, and green indicates neutralization capacity. *** denotes multiple variants. (**b**) Representative binding curve for mAb G4 competing with pooled serially diluted sera from mice immunized with SS-L2-LuSNP. $R_0$ represents the binding signal of G4 in the absence of serum, and Rc represents the binding signal of G4 tested with sera.

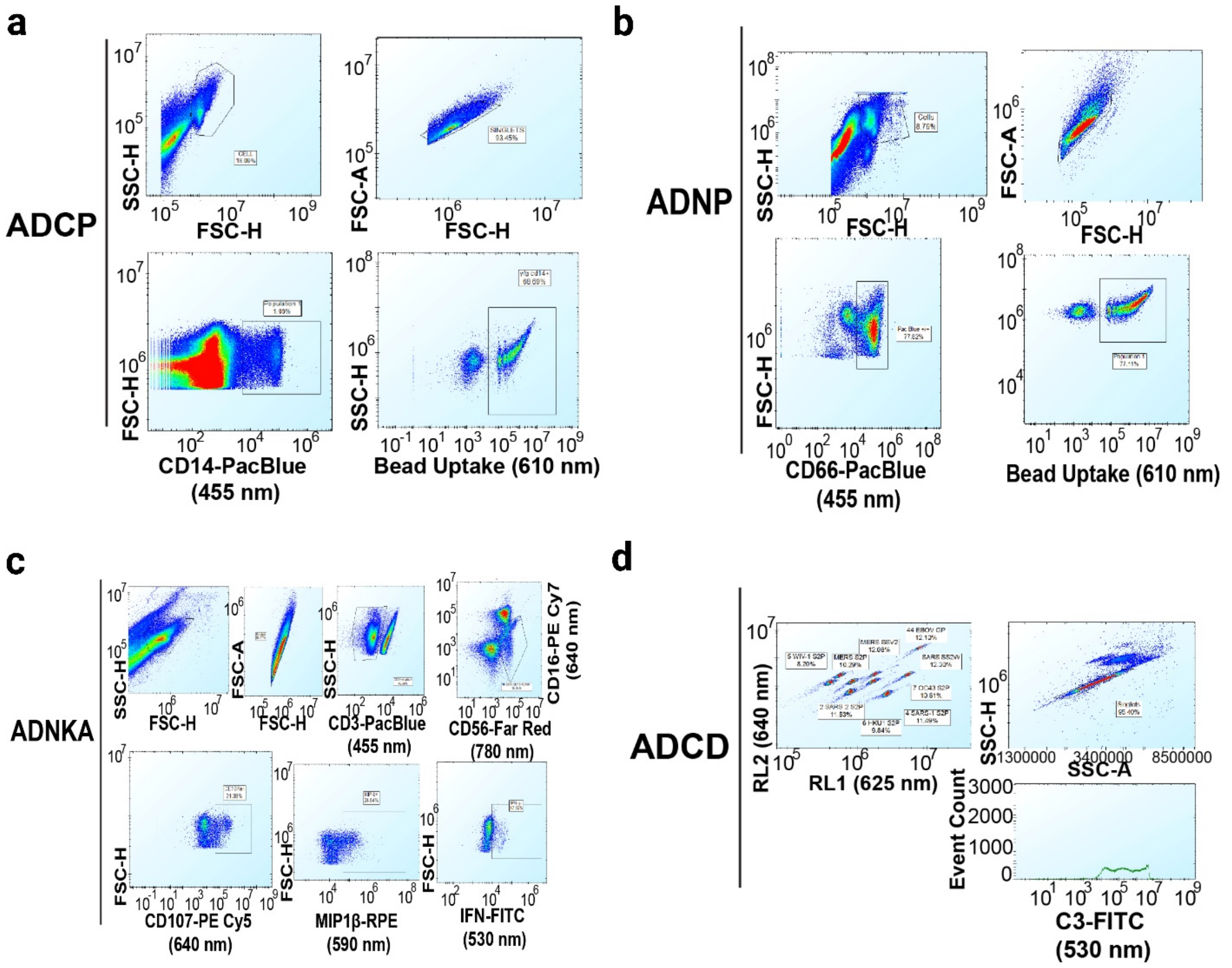


**Supplementary Fig. 2: Gating strategy for Luminex-based and Ab Fc-effector function assays, related to Fig. 5.** (**a**) Gating for antibody-dependent cellular phagocytosis by monocytes (ADCP) showing CD14+ monocytes and opsinophagocytosis. (**b**) Gating for antibody-dependent neutrophil phagocytosis (ADNP) showing CD66+ neutrophils and opsinophagocytosis (**c**) Gating for antibody-dependent natural killer cell activation (ADNKA) showing CD107a and MIP-1β expression on CD3-/CD56+/CD16- cells. (**d**) Gating for antibody-dependent complement deposition (ADCD) on CoV proteins on individual bead regions as measured by anti C3-FITC.

**Supplementary Table 1:** Conventional and *de novo* designed (unconventional) linkers.

| Linker Type | Sequence | References |
|---|---|---|
| **Short Flexible (L0)** | GG; GS; GSG; SGG; GGGGS | 1-19 |
| **Long Flexible (L1)** | 8(GS); 12(GS); 16(GS); 2(GGGGS); 4(GGS);3(GGGGS) | 11,20-25 |
| **Rigid Helical (L2)** | GGGGSAEAAAKEAAAKE EAAAKAGGGGS | This Study |
| **Long Flexible (L6)** | GGGSGGGGSGGGGSLSK | 26 |

**Supplementary Table 2.** Cryo-EM data collection and processing parameters, related to Fig. 2e,f.

| Cryo-EM Data Collection and Processing Statistics | | | |
|---|---|---|---|
| **EM Data Collection** | SS-L2-FerritinNP | SS-L2-LuSNP | SS-L2-153-50NP |
| Microscope | Talos Arctica | Talos Arctica | Talos Arctica |
| Software | SerialEM | SerialEM | SerialEM |
| Voltage(kV) | 200 | 200 | 200 |
| Detector | Gatan K3 | Gatan K3 | Gatan K3 |
| Magnification(Norminal) | 36,000 | 36,000 | 36,000 |
| Electron exposure(e-/ Å$^2$) | 50.12 | 60.57 | 57.39 |
| Defocus range (μm) | -0.6-2.4 | -0.8-2.0 | 0.9-2.4 |
| Pixel size(Å) | 1.1 | 1.1 | 1.1 |
| Flux (e-/pix/ Å) | 13.06 | 15.9 | 13.88 |
| Exposure time(seconds) | 4.89 | 4 | 5 |
| Total Frames | 49 | 46 | 50 |
| Micrographs collected | 1333 | 820 | 1939 |

| 3D Reconstruction Statistics | | | |
|---|---|---|---|
| Software | CryoSPARC v4.6.2 | CryoSPARC v4.6.2 | CryoSPARC v4.6.2 |
| Symmetry imposed | O | T | I |
| Box size(pix) | 450 | 450 | 450 |
| Initial particle image no | 1333 | 820 | 1939 |
| Final particle image no | 1300 | 800 | 1900 |
| Map sharpening B-factor | -149 | -34 | -100 |
| FSC threshold | 0.143 | 0.143 | 0.143 |
| Unmasked resolution at 0.143FSC(Å) | 7.3 | 7.9 | 6.2 |
| Masked resolution at 0.143FSC(Å) | 6.2 | 4.1 | 4.4 |
| Map resolution(Å) | 6.25 | 4.22 | 4.42 |

**Supplementary Table 3:** Sequences of designed immunogens and expression constructs reported in this study.

| Construct Name | Amino Acid Sequence |
|---|---|
| **Color Key** | **Secretion Signal Sequence**,MERS Stabilized S-2P, MERS-CoV Stabilized Stem(SS), Display Linker(L2), Display Linker(L6), Foldon, Streptag II, His-8-tag, FerritinNP, Lumazine Synthase (LuS)NP, I53-50A.1NT1, I53-50B.4PT1, 3C Protease Cleavage |
| **MERS S-2P** | **MIHSVFLLMFLLTPTES**YVDVGPDSVKSACIEVDIQQTFFDKTWPRPIDVSKADGIIYPQGRTYSNITITYQGLFPYQGDHGDMYVYSAGHATGTTPQKLFVANYSQDVKQFANGFVVRIGAAANSTGTVIISPSTSATIRKIYPAFMLGSSVGNFSDGKMGRFFNHTLVLLPDGCGTLLRAFYCILEPRSGNHCPAGNSYTSFATYHTPATDCSDGNYNRNASLNSFKEYFNLRNCTFMYTYNITEDEILEWFGITQTAQGVHLFSSRYVDLYGGNMFQFATLPVYDTIKYYSIIPHSIRSIQSDRKAWAAFYVYKLQPLTFLLDFSVDGYIRRAIDCGFNDLSQLHCSYESFDVESGVYSVSSFEAKPSGSVVEQAEGVECDFSPLLSGTPPQVYNFKRLVFTNCNYNLTKLLSLFSVNDFTCSQISPAAIASNCYSSLILDYFSYPLSMKSDLSVSSAGPISQFNYKQSFSNPTCLILATVPHNLTTITKPLKYSYINKCSRFLSDDRTEVPQLVNANQYSPCVSIVPSTVWEDGDYYRKQLSPLEGGGWLVASGSTVAMTEQLQMGFGITVQYGTDTNSVCPKLEFANDTKIASQLGNCVEYSLYGVSGRGVFQNCTAVGVRQQRFVYDAYQNLVGYYSDDGNYYCLRACVSVPVSVIYDKETKTHATLFGSVACEHISSTMSQYSRSTRSMLKRRDSTYGPLQTPVGCVLGLVNSSLFVEDCKLPLGQSLCALPDTPSTLTPASVGSVPGEMRLASIAFNHPIQVDQLNSSYFKLSIPTNFSFGVTQEYIQTTIQKVTVDCKQYVCNGFQKCEQLLREYGQFCSKINQALHGANLRQDDSVRNLFASVKSSQSSPIIPGFGGDFNLTLLEPVSISTGSRSARSAIEDLLFDKVTIADPGYMQGYDDCMQQGPASARDLICAQYVAGYKVLPPLMDVNMEAAYTSSLLGSIAGVGWTAGLSSFAAIPFAQSIFYRLNGVGITQQVLSENQKLIANKFNQALGAMQTGFTTTNEAFHKVQDAVNNNAQALSKLASELSNTFGAISASIGDIIQRLDPPEQDAQIDRLINGRLTTLNAFVAQQLVRSESAALSAQLAKDKVNECVKAQSKRSGFCGQGTHIVSFVVNAPNGLYFMHVGYYPSNHIEVVSAYGLCDAANPTNCIAPVNGYFIKTNNTRIVDEWSYTGSSFYAPEPITSLNTKYVAPQVTYQNISTNLPPPLLGNSTGIDFQDELDEFFKNVSTSIPNFGSLTQINTTLLDLTYEMLSLQQVVKALNESYIDLKELGNYTYGSGYIPEAPRDGQAYVRKDGEWVLLSTFLGRSLEVLFQGPHHHHHHHHSAWSHPQFEKGGGSGGGGSGGSAWSHPQFEK |
| **MERS S-2P-L2-FerritinNP** | **MIHSVFLLMFLLTPTES**YVDVGPDSVKSACIEVDIQQTFFDKTWPRPIDVSKADGIIYPQGRTYSNITITYQGLFPYQGDHGDMYVYSAGHATGTTPQKLFVANYSQDVKQFANGFVVRIGAAANSTGTVIISPSTSATIRKIYPAFMLGSSVGNFSDGKMGRFFNHTLVLLPDGCGTLLRAFYCILEPRSGNHCPAGNSYTSFATYHTPATDCSDGNYNRNASLNSFKEYFNLRNCTFMYTYNITEDEILEWFGITQTAQGVHLFSSRYVDLYGGNMFQFATLPVYDTIKYYSIIPHSIRSIQSDRKAWAAFYVYKLQPLTFLLDFSVDGYIRRAIDCGFNDLSQLHCSYESFDVESGVYSVSSFEAKPSGSVVEQAEGVECDFSPLLSGTPPQVYNFKRLVFTNCNYNLTKLLSLFSVNDFTCSQISPAAIASNCYSSLILDYFSYPLSMKSDLSVSSAGPISQFNYKQSFSNPTCLILATVPHNLTTITKPLKYSYINKCSRFLSDDRTEVPQLVNANQYSPCVSIVPSTVWEDGDYYRKQLSPLEGGGWLVASGSTVAMTEQLQMGFGITVQYGTDTNSVCPKLEFANDTKIASQLGNCVEYSLYGVSGRGVFQNCTAVGVRQQRFVYDAYQNLVGYYSDDGNYYCLRACVSVPVSVIYDKETKTHATLFGSVACEHISSTMSQYSRSTRSMLKRRDSTYGPLQTPVGCVLGLVNSSLFVEDCKLPLGQSLCALPDTPSTLTPASVGSVPGEMRLASIAFNHPIQVDQLNSSYFKLSIPTNFSFGVTQEYIQTTIQKVTVDCKQYVCNGFQKCEQLLREYGQFCSKINQALHGANLRQDDSVRNLFASVKSSQSSPIIPGFGGDFNLTLLEPV |

| | |
|---|---|
| | SISTGSRSARSAIEDLLFDKVTIADPGYMQGYDDCMQQGPASARDLICAQYVAGYKVLPPLMDVNMEAAYTSSLLGSIAGVGWTAGL<br>SSFAAIPFAQSIFYRLNGVGITQQVLSENQKLIANKFNQALGAMQTGFTTTNEAFHKVQDAVNNNAQALSKLASELSNTFGAISASIGD<br>IIQRLDPPEQDAQIDRLINGRLTTLNAFVAQQLVRSESAALSAQLAKDKVNECVKAQSKRSGFCGQGTHIVSFVVNAPNGLYFMHVG<br>YYPSNHIEVVSAYGLCDAANPTNCIAPVNGYFIKTNNTRIVDEWSYTGSSFYAPEPITSLNTKYVAPQVTYQNISTNLPPPLLGNSTGI<br>DFQDELDEFFKNVSTSIPN F GGGGS AEAAAKEAAAKEAAAKA<br>GGGGSMSSQIRQNYSTDVEAAVNSLVNLYLQASYTYLSLGFYFDRDDVALEGVSHFFRELAEEKREGYERLLKMQNQRGGRALFQ<br>DIKKPAEDEWGKTPDAMKAAMALEKKLNQALLDLHALGSARTDPHLCDFLETHFLDEEVKLIKKMGDHLTNLHRLGGPEAGLGEYLF<br>ERLTLKHD |
| **MERS S-2P-L6-FerritinNP** | **MIHSVFLLMFLLTPTES**YVDVGPDSVKSACIEVDIQQTFFDKTWPRPIDVSKADGIIYPQGRTYSNITITYQGLFPYQGDHGDMYVYS<br>AGHATGTTPQKLFVANYSQDVKQFANGFVVRIGAAANSTGTVIISPSTSATIRKIYPAFMLGSSVGNFSDGKMGRFFNHTLVLLPDGC<br>GTLLRAFYCILEPRSGNHCPAGNSYTSFATYHTPATDCSDGNYNRNASLNSFKEYFNLRNCTFMYTYNITEDEILEWFGITQTAQGVH<br>LFSSRYVDLYGGNMFQFATLPVYDTIKYYSIIPHSIRSIQSDRKAWAAFYVYKLQPLTFLLDFSVDGYIRRAIDCGFNDLSQLHCSYESF<br>DVESGVYSVSSFEAKPSGSVVEQAEGVECDFSPLLSGTPPQVYNFKRLVFTNCNYNLTKLLSLFSVNDFTCSQISPAAIASNCYSSLI<br>LDYFSYPLSMKSDLSVSSAGPISQFNYKQSFSNPTCLILATVPHNLTTITKPLKYSYINKCSRFLSDDRTEVPQLVNANQYSPCVSIVP<br>STVWEDGDYYRKQLSPLEGGGWLVASGSTVAMTEQLQMGFGITVQYGTDTNSVCPKLEFANDTKIASQLGNCVEYSLYGVSGRGV<br>FQNCTAVGVRQQRFVYDAYQNLVGYYSDDGNYYCLRACVSVPVSVIYDKETKTHATLFGSVACEHISSTMSQYSRSTRSMLKRRDS<br>TYGPLQTPVGCVLGLVNSSLFVEDCKLPLGQSLCALPDTPSTLTPASVGSVPGEMRLASIAFNHPIQVDQLNSSYFKLSIPTNFSFGV<br>TQEYIQTTIQKVTVDCKQYVCNGFQKCEQLLREYGQFCSKINQALHGANLRQDDSVRNLFASVKSSQSSPIIPGFGGDFNLTLLEPV<br>SISTGSRSARSAIEDLLFDKVTIADPGYMQGYDDCMQQGPASARDLICAQYVAGYKVLPPLMDVNMEAAYTSSLLGSIAGVGWTAGL<br>SSFAAIPFAQSIFYRLNGVGITQQVLSENQKLIANKFNQALGAMQTGFTTTNEAFHKVQDAVNNNAQALSKLASELSNTFGAISASIGD<br>IIQRLDPPEQDAQIDRLINGRLTTLNAFVAQQLVRSESAALSAQLAKDKVNECVKAQSKRSGFCGQGTHIVSFVVNAPNGLYFMHVG<br>YYPSNHIEVVSAYGLCDAANPTNCIAPVNGYFIKTNNTRIVDEWSYTGSSFYAPEPITSLNTKYVAPQVTYQNISTNLPPPLLGNSTGI<br>DFQDELDEFFKNVSTSIPN FGS<br>GGGSGGGGSGGGGSGLSKMSSQIRQNYSTDVEAAVNSLVNLYLQASYTYLSLGFYFDRDDVALEGVSHFFRELAEEKREGYERLL<br>KMQNQRGGRALFQDIKKPAEDEWGKTPDAMKAAMALEKKLNQALLDLHALGSARTDPHLCDFLETHFLDEEVKLIKKMGDHLTNLH<br>RLGGPEAGLGEYLFERLTLKHD |
| **MERS S-2P-L2-LuSNP** | **MIHSVFLLMFLLTPTES**YVDVGPDSVKSACIEVDIQQTFFDKTWPRPIDVSKADGIIYPQGRTYSNITITYQGLFPYQGDHGDMYVYS<br>AGHATGTTPQKLFVANYSQDVKQFANGFVVRIGAAANSTGTVIISPSTSATIRKIYPAFMLGSSVGNFSDGKMGRFFNHTLVLLPDGC<br>GTLLRAFYCILEPRSGNHCPAGNSYTSFATYHTPATDCSDGNYNRNASLNSFKEYFNLRNCTFMYTYNITEDEILEWFGITQTAQGVH<br>LFSSRYVDLYGGNMFQFATLPVYDTIKYYSIIPHSIRSIQSDRKAWAAFYVYKLQPLTFLLDFSVDGYIRRAIDCGFNDLSQLHCSYESF<br>DVESGVYSVSSFEAKPSGSVVEQAEGVECDFSPLLSGTPPQVYNFKRLVFTNCNYNLTKLLSLFSVNDFTCSQISPAAIASNCYSSLI<br>LDYFSYPLSMKSDLSVSSAGPISQFNYKQSFSNPTCLILATVPHNLTTITKPLKYSYINKCSRFLSDDRTEVPQLVNANQYSPCVSIVP<br>STVWEDGDYYRKQLSPLEGGGWLVASGSTVAMTEQLQMGFGITVQYGTDTNSVCPKLEFANDTKIASQLGNCVEYSLYGVSGRGV<br>FQNCTAVGVRQQRFVYDAYQNLVGYYSDDGNYYCLRACVSVPVSVIYDKETKTHATLFGSVACEHISSTMSQYSRSTRSMLKRRDS |

| | |
|---|---|
| | TYGPLQTPVGCVLGLVNSSLFVEDCKLPLGQSLCALPDTPSTLTPASVGSVPGEMRLASIAFNHPIQVDQLNSSYFKLSIPTNFSFGV<br>TQEYIQTTIQKVTVDCKQYVCNGFQKCEQLLREYGQFCSKINQALHGANLRQDDSVRNLFASVKSSQSSPIIPGFGGDFNLTLLEPV<br>SISTGSRSARSAIEDLLFDKVTIADPGYMQGYDDCMQQGPASARDLICAQYVAGYKVLPPLMDVNMEAAYTSSLLGSIAGVGWTAGL<br>SSFAAIPFAQSIFYRLNGVGITQQVLSENQKLIANKFNQALGAMQTGFTTTNEAFHKVQDAVNNNAQALSKLASELSNTFGAISASIGD<br>IIQRLDPPEQDAQIDRLINGRLTTLNAFVAQQLVRSESAALSAQLAKDKVNECVKAQSKRSGFCGQGTHIVSFVVNAPNGLYFMHVG<br>YYPSNHIEVVSAYGLCDAANPTNCIAPVNGYFIKTNNTRIVDEWSYTGSSFYAPEPITSLNTKYVAPQVTYQNISTNLPPPLLGNSTGI<br>DFQDELDEFFKNVSTSIPN FGGGGS AEAAAKEAAAKEAAAKA GGGGS<br>MQIYEGKLTAEGLRFGIVASRFNHALVDRLVEGAIDCIVRHGGREEDITLVRVPGSWEIPVAAGELARKEDIDAVIAIGVLIRGATPHFDY<br>IASEVSKGLANLSLELRKPITFGVITADTLEQAIERAGTKHGNKGWEAALSAIEMANLFKSLR |
| **MERS S-2P-L6-LuSNP** | **MIHSVFLLMFLLTPTES**YVDVGPDSVKSACIEVDIQQTFFDKTWPRPIDVSKADGIIYPQGRTYSNITITYQGLFPYQGDHGDMYVYS<br>AGHATGTTPQKLFVANYSQDVKQFANGFVVRIGAAANSTGTVIISPSTSATIRKIYPAFMLGSSVGNFSDGKMGRFFNHTLVLLPDGC<br>GTLLRAFYCILEPRSGNHCPAGNSYTSFATYHTPATDCSDGNYNRNASLNSFKEYFNLRNCTFMYTYNITEDEILEWFGITQTAQGVH<br>LFSSRYVDLYGGNMFQFATLPVYDTIKYYSIIPHSIRSIQSDRKAWAAFYVYKLQPLTFLLDFSVDGYIRRAIDCGFNDLSQLHCSYESF<br>DVESGVYSVSSFEAKPSGSVVEQAEGVECDFSPLLSGTPPQVYNFKRLVFTNCNYNLTKLLSLFSVNDFTCSQISPAAIASNCYSSLI<br>LDYFSYPLSMKSDLSVSSAGPISQFNYKQSFSNPTCLILATVPHNLTTITKPLKYSYINKCSRFLSDDRTEVPQLVNANQYSPCVSIVP<br>STVWEDGDYYRKQLSPLEGGGWLVASGSTVAMTEQLQMGFGITVQYGTDTNSVCPKLEFANDTKIASQLGNCVEYSLYGVSGRGV<br>FQNCTAVGVRQQRFVYDAYQNLVGYYSDDGNYYCLRACVSVPVSVIYDKETKTHATLFGSVACEHISSTMSQYSRSTRSMLKRRDS<br>TYGPLQTPVGCVLGLVNSSLFVEDCKLPLGQSLCALPDTPSTLTPASVGSVPGEMRLASIAFNHPIQVDQLNSSYFKLSIPTNFSFGV<br>TQEYIQTTIQKVTVDCKQYVCNGFQKCEQLLREYGQFCSKINQALHGANLRQDDSVRNLFASVKSSQSSPIIPGFGGDFNLTLLEPV<br>SISTGSRSARSAIEDLLFDKVTIADPGYMQGYDDCMQQGPASARDLICAQYVAGYKVLPPLMDVNMEAAYTSSLLGSIAGVGWTAGL<br>SSFAAIPFAQSIFYRLNGVGITQQVLSENQKLIANKFNQALGAMQTGFTTTNEAFHKVQDAVNNNAQALSKLASELSNTFGAISASIGD<br>IIQRLDPPEQDAQIDRLINGRLTTLNAFVAQQLVRSESAALSAQLAKDKVNECVKAQSKRSGFCGQGTHIVSFVVNAPNGLYFMHVG<br>YYPSNHIEVVSAYGLCDAANPTNCIAPVNGYFIKTNNTRIVDEWSYTGSSFYAPEPITSLNTKYVAPQVTYQNISTNLPPPLLGNSTGI<br>DFQDELDEFFKNVSTSIPN FGS GGGSGGGGSGGGGSGLSK<br>MQIYEGKLTAEGLRFGIVASRFNHALVDRLVEGAIDCIVRHGGREEDITLVRVPGSWEIPVAAGELARKEDIDAVIAIGVLIRGATPHFDY<br>IASEVSKGLANLSLELRKPITFGVITADTLEQAIERAGTKHGNKGWEAALSAIEMANLFKSLR |
| **SS** | **MRPTWAWWLFLVLLLALWAPARG**QAFNHPIQVDQLNSSYFKLSIPTNFSFGVTQEYIQTTIQKVCVDCKQYVCNGFQKCEQLLREY<br>GQFCSKINQALHGCNLRQDDSVRNLFASVKSSQSSPIIPGFGGDFNLTLLEPVSISTGSRSARSAIEDLLFDKCTIADPGYMQGYDDC<br>MQQGPASARDLICAQYVAGYCVLPPLMDVNMEAAYTSSLLGSIAGSGWTAGLSSFAAIPFAQMIFYRLNGIGITQQVLSENQKLIANK<br>FNQALGAMQTGFTTTNEAFHKCQDAVNNNAQALSKLASELSNTFGAISASIGDIIQRLDPPEQDAQIDRLINGRLTTLNAFVAQQLVR<br>CEEAAQSAQLAKDKVNECVKAQSKRSGFCGQGTHIVSFVVNAPYGLYFMHVGYYPSNHIEVVSAYGLCDAANPTNCIAPVNGYFIK<br>TNNTRIVDEWSYTGSSFYAPEPITSLNTKYVAPQVTYQNISTNLPPPLLGNSTGIDFQDELDEFFKNVSTSIPNFGSLTQINTTLLDLTY<br>EMLSLQQVVKALNESYIDLKELGNYTYGSGYIPEAPRDGQAYVRKDGEWVLLSTFLGGRLEVLFQGPGGYIPEAPRDGQAYVRKDG<br>EWVLLSTFLGHHHHHHHHSAWSHPQFEK |

| | |
|---|---|
| **SS-L2-FerritinNP** | **MRPTWAWWLFLVLLLALWAPARG**QAFNHPIQVDQLNSSYFKLSIPTNFSFGVTQEYIQTTIQKVCVDCKQYVCNGFQKCEQLLREY<br>GQFCSKINQALHGCNLRQDDSVRNLFASVKSSQSSPIIPGFGGDFNLTLLEPVSISTGSRSARSAIEDLLFDKCTIADPGYMQGYDDC<br>MQQGPASARDLICAQYVAGYCVLPPLMDVNMEAAYTSSLLGSIAGSGWTAGLSSFAAIPFAQMIFYRLNGIGITQQVLSENQKLIANK<br>FNQALGAMQTGFTTTNEAFHKCQDAVNNNAQALSKLASELSNTFGAISASIGDIIQRLDPPEQDAQIDRLINGRLTTLNAFVAQQLVR<br>CEEAAQSAQLAKDKVNECVKAQSKRSGFCGQGTHIVSFVVNAPYGLYFMHVGYYPSNHIEVVSAYGLCDAANPTNCIAPVNGYFIK<br>TNNTRIVDEWSYTGSSFYAPEPITSLNTKYVAPQVTYQNISTNLPPPLLGNSTGIDFQDELDEFFKNVSTSIPNFGSLTQINTTLLDLTY<br>EMLSLQQVVKALNESYIDLKELGNYTYGSGYIPEAPRDGQAYVRKDGEWVLLSTFLG**GGGGSAEAAAKEAAAKEAAAKAGGGGS**<br>MSSQIRQNYSTDVEAAVNSLVNLYLQASYTYLSLGFYFDRDDVALEGVSHFFRELAEEKREGYERLLKMQNQRGGRALFQDIKKPA<br>EDEWGKTPDAMKAAMALEKKLNQALLDLHALGSARTDPHLCDFLETHFLDEEVKLIKKMGDHLTNLHRLGGPEAGLGEYLFERLTLK<br>HD |
| **SS-L6-FerritinNP** | **MRPTWAWWLFLVLLLALWAPARG**QAFNHPIQVDQLNSSYFKLSIPTNFSFGVTQEYIQTTIQKVCVDCKQYVCNGFQKCEQLLREY<br>GQFCSKINQALHGCNLRQDDSVRNLFASVKSSQSSPIIPGFGGDFNLTLLEPVSISTGSRSARSAIEDLLFDKCTIADPGYMQGYDDC<br>MQQGPASARDLICAQYVAGYCVLPPLMDVNMEAAYTSSLLGSIAGSGWTAGLSSFAAIPFAQMIFYRLNGIGITQQVLSENQKLIANK<br>FNQALGAMQTGFTTTNEAFHKCQDAVNNNAQALSKLASELSNTFGAISASIGDIIQRLDPPEQDAQIDRLINGRLTTLNAFVAQQLVR<br>CEEAAQSAQLAKDKVNECVKAQSKRSGFCGQGTHIVSFVVNAPYGLYFMHVGYYPSNHIEVVSAYGLCDAANPTNCIAPVNGYFIK<br>TNNTRIVDEWSYTGSSFYAPEPITSLNTKYVAPQVTYQNISTNLPPPLLGNSTGIDFQDELDEFFKNVSTSIPNFGSLTQINTTLLDLTY<br>EMLSLQQVVKALNESYIDLKELGNYTYGSGYIPEAPRDGQAYVRKDGEWVLLSTFLG**GGGSGGGGSGGGGSGLSK**MSSQIRQNY<br>STDVEAAVNSLVNLYLQASYTYLSLGFYFDRDDVALEGVSHFFRELAEEKREGYERLLKMQNQRGGRALFQDIKKPAEDEWGKTPD<br>AMKAAMALEKKLNQALLDLHALGSARTDPHLCDFLETHFLDEEVKLIKKMGDHLTNLHRLGGPEAGLGEYLFERLTLKHD |
| **SS-L2-LuSNP** | **MRPTWAWWLFLVLLLALWAPARG**QAFNHPIQVDQLNSSYFKLSIPTNFSFGVTQEYIQTTIQKVCVDCKQYVCNGFQKCEQLLREY<br>GQFCSKINQALHGCNLRQDDSVRNLFASVKSSQSSPIIPGFGGDFNLTLLEPVSISTGSRSARSAIEDLLFDKCTIADPGYMQGYDDC<br>MQQGPASARDLICAQYVAGYCVLPPLMDVNMEAAYTSSLLGSIAGSGWTAGLSSFAAIPFAQMIFYRLNGIGITQQVLSENQKLIANK<br>FNQALGAMQTGFTTTNEAFHKCQDAVNNNAQALSKLASELSNTFGAISASIGDIIQRLDPPEQDAQIDRLINGRLTTLNAFVAQQLVR<br>CEEAAQSAQLAKDKVNECVKAQSKRSGFCGQGTHIVSFVVNAPYGLYFMHVGYYPSNHIEVVSAYGLCDAANPTNCIAPVNGYFIK<br>TNNTRIVDEWSYTGSSFYAPEPITSLNTKYVAPQVTYQNISTNLPPPLLGNSTGIDFQDELDEFFKNVSTSIPNFGSLTQINTTLLDLTY<br>EMLSLQQVVKALNESYIDLKELGNYTYGSGYIPEAPRDGQAYVRKDGEWVLLSTFLG**GGGGSAEAAAKEAAAKEAAAKAGGGGS**<br>MQIYEGKLTAEGLRFGIVASRFNHALVDRLVEGAIDCIVRHGGREEDITLVRVPGSWEIPVAAGELARKEDIDAVIAIGVLIRGATPHFDY<br>IASEVSKGLANLSLELRKPITFGVITADTLEQAIERAGTKHGNKGWEAALSAIEMANLFKSLR |
| **SS-L6-LuSNP** | **MRPTWAWWLFLVLLLALWAPARG**QAFNHPIQVDQLNSSYFKLSIPTNFSFGVTQEYIQTTIQKVCVDCKQYVCNGFQKCEQLLREY<br>GQFCSKINQALHGCNLRQDDSVRNLFASVKSSQSSPIIPGFGGDFNLTLLEPVSISTGSRSARSAIEDLLFDKCTIADPGYMQGYDDC<br>MQQGPASARDLICAQYVAGYCVLPPLMDVNMEAAYTSSLLGSIAGSGWTAGLSSFAAIPFAQMIFYRLNGIGITQQVLSENQKLIANK<br>FNQALGAMQTGFTTTNEAFHKCQDAVNNNAQALSKLASELSNTFGAISASIGDIIQRLDPPEQDAQIDRLINGRLTTLNAFVAQQLVR<br>CEEAAQSAQLAKDKVNECVKAQSKRSGFCGQGTHIVSFVVNAPYGLYFMHVGYYPSNHIEVVSAYGLCDAANPTNCIAPVNGYFIK<br>TNNTRIVDEWSYTGSSFYAPEPITSLNTKYVAPQVTYQNISTNLPPPLLGNSTGIDFQDELDEFFKNVSTSIPNFGSLTQINTTLLDLTY |

| | |
|---|---|
| | EMLSLQQVVKALNESYIDLKELGNYTYGSGYIPEAPRDGQAYVRKDGEWVLLSTFLG**GGGSGGGGSGGGGSGLSK**MQIYEGKLTA<br>EGLRFGIVASRFNHALVDRLVEGAIDCIVRHGGREEDITLVRVPGSWEIPVAAGELARKEDIDAVIAIGVLIRGATPHFDYIASEVSKGLA<br>NLSLELRKPITFGVITADTLEQAIERAGTKHGNKGWEAALSAIEMANLFKSLR |
| **SS-L2-I53-50A.1NT1** | **MRPTWAWWLFLVLLLALWAPARG**QAFNHPIQVDQLNSSYFKLSIPTNFSFGVTQEYIQTTIQKVCVDCKQYVCNGFQKCEQLLREY<br>GQFCSKINQALHGCNLRQDDSVRNLFASVKSSQSSPIIPGFGGDFNLTLLEPVSISTGSRSARSAIEDLLFDKCTIADPGYMQGYDDC<br>MQQGPASARDLICAQYVAGYCVLPPLMDVNMEAAYTSSLLGSIAGSGWTAGLSSFAAIPFAQMIFYRLNGIGITQQVLSENQKLIANK<br>FNQALGAMQTGFTTTNEAFHKCQDAVNNNAQALSKLASELSNTFGAISASIGDIIQRLDPPEQDAQIDRLINGRLTTLNAFVAQQLVR<br>CEEAAQSAQLAKDKVNECVKAQSKRSGFCGQGTHIVSFVVNAPYGLYFMHVGYYPSNHIEVVSAYGLCDAANPTNCIAPVNGYFIK<br>TNNTRIVDEWSYTGSSFYAPEPITSLNTKYVAPQVTYQNISTNLPPPLLGNSTGIDFQDELDEFFKNVSTSIPNFGSLTQINTTLLDLTY<br>EMLSLQQVVKALNESYIDLKELGNYTYGSGYIPEAPRDGQAYVRKDGEWVLLSTFLG**GGGGSAEAAAKEAAAKEAAAKAGGGGS**<br>MEELFKKHKIVAVLRANSVEEAIEKAVAVFAGGVHLIEITFTVPDADTVIKALSVLKEKGAIIGAGTVTSVEQCRKAVESGAEFIVSPHL<br>DEEISQFCKEKGVFYMPGVMTPTELVKAMKLGHDILKLFPGEVVGPEFVKAMKGPFPNVKFVPTGGVDLDNVCEWFDAGVLAVGV<br>GDALVEGDPDEVREKAKEFVEKIRGCTEGSLEWSHPQFEKGSGHHHHHHHH |
| **SS-L6-I53-50A.1NT1** | **MRPTWAWWLFLVLLLALWAPARG**QAFNHPIQVDQLNSSYFKLSIPTNFSFGVTQEYIQTTIQKVCVDCKQYVCNGFQKCEQLLREY<br>GQFCSKINQALHGCNLRQDDSVRNLFASVKSSQSSPIIPGFGGDFNLTLLEPVSISTGSRSARSAIEDLLFDKCTIADPGYMQGYDDC<br>MQQGPASARDLICAQYVAGYCVLPPLMDVNMEAAYTSSLLGSIAGSGWTAGLSSFAAIPFAQMIFYRLNGIGITQQVLSENQKLIANK<br>FNQALGAMQTGFTTTNEAFHKCQDAVNNNAQALSKLASELSNTFGAISASIGDIIQRLDPPEQDAQIDRLINGRLTTLNAFVAQQLVR<br>CEEAAQSAQLAKDKVNECVKAQSKRSGFCGQGTHIVSFVVNAPYGLYFMHVGYYPSNHIEVVSAYGLCDAANPTNCIAPVNGYFIK<br>TNNTRIVDEWSYTGSSFYAPEPITSLNTKYVAPQVTYQNISTNLPPPLLGNSTGIDFQDELDEFFKNVSTSIPNFGSLTQINTTLLDLTY<br>EMLSLQQVVKALNESYIDLKELGNYTYGSGYIPEAPRDGQAYVRKDGEWVLLSTFLG**GGGSGGGGSGGGGSGLSK**MEELFKKHKI<br>VAVLRANSVEEAIEKAVAVFAGGVHLIEITFTVPDADTVIKALSVLKEKGAIIGAGTVTSVEQCRKAVESGAEFIVSPHLDEEISQFCKE<br>KGVFYMPGVMTPTELVKAMKLGHDILKLFPGEVVGPEFVKAMKGPFPNVKFVPTGGVDLDNVCEWFDAGVLAVGVGDALVEGDP<br>DEVREKAKEFVEKIRGCTEGSLEWSHPQFEKGSGHHHHHHHH |
| **I53-50A.1NT1** | MKMEELFKKHKIVAVLRANSVEEAIEKAVAVFAGGVHLIEITFTVPDADTVIKALSVLKEKGAIIGAGTVTSVEQCRKAVESGAEFIVSP<br>HLDEEISQFCKEKGVFYMPGVMTPTELVKAMKLGHDILKLFPGEVVGPEFVKAMKGPFPNVKFVPTGGVDLDNVCEWFDAGVLAV<br>GVGDALVEG DPDEVREKAKEFVEKIRGCTEGSLEHHHHHHHH |
| **I53-50B.4PT1** | MNQHSHKDHETVRIAVVRARWHAEIVDACVSAFEAAMRDIGGDRFAVDVFDVPGAYEIPLHARTLAETGRYGAVLGTAFVVNGGIY<br>RHEFVASAVINGMMNVQLNTGVPVLSAVLTPHNYDKSKAHTLLFLALFAVKGMEAARACVEILAAREKIAAGSLEHHHHHHHH |

### References, related to Supplementary Table 1


1 Brinkkemper, M. *et al.* Co-display of diverse spike proteins on nanoparticles broadens sarbecovirus neutralizing antibody responses. *iScience* **25**, 105649-105649 (2022). https://doi.org/10.1016/j.isci.2022.105649

2 Joyce, M. G. *et al.* SARS-CoV-2 ferritin nanoparticle vaccines elicit broad SARS coronavirus immunogenicity. *Cell Reports* **37**, 110143 (2021). https://doi.org/10.1016/j.celrep.2021.110143

3 Hutchinson, G. B. *et al.* Nanoparticle display of prefusion coronavirus spike elicits S1-focused cross-reactive antibody response against diverse coronavirus subgenera. *Nature Communications* **14** (2023). https://doi.org/10.1038/s41467-023-41661-4

4 Antanasijevic, A. *et al.* Structural and functional evaluation of de novo-designed, two-component nanoparticle carriers for HIV Env trimer immunogens. *PLOS Pathogens* **16**, e1008665 (2020). https://doi.org/10.1371/journal.ppat.1008665

5 Boyoglu-Barnum, S. *et al.* Quadrivalent influenza nanoparticle vaccines induce broad protection. *Nature* **592**, 623-628 (2021). https://doi.org/10.1038/s41586-021-03365-x

6 Kraft, J. C. *et al.* Antigen- and scaffold-specific antibody responses to protein nanoparticle immunogens. *Cell Reports Medicine* **3**, 100780 (2022). https://doi.org/10.1016/j.xcrm.2022.100780

7 Kanekiyo, M. *et al.* Mosaic nanoparticle display of diverse influenza virus hemagglutinins elicits broad B cell responses. *Nature Immunology* **20**, 362-372 (2019). https://doi.org/10.1038/s41590-018-0305-x

8 Ueda, G. *et al.* (Cold Spring Harbor Laboratory, 2020).

9 Brouwer, P. J. M. *et al.* Lassa virus glycoprotein nanoparticles elicit neutralizing antibody responses and protection. *Cell Host & Microbe* **30**, 1759-1772.e1712 (2022). https://doi.org/10.1016/j.chom.2022.10.018

10 Hu, Z. *et al.* Nanoparticle vaccine based on the pre-fusion F glycoprotein of respiratory syncytial virus elicits robust protective immune responses. *Journal of Virology* **99** (2025). https://doi.org/10.1128/jvi.00903-25

11 Zhang, Y.-N. *et al.* Single-component multilayered self-assembling protein nanoparticles presenting glycan-trimmed uncleaved prefusion optimized envelope trimers as HIV-1 vaccine candidates. *Nature Communications* **14** (2023). https://doi.org/10.1038/s41467-023-37742-z

12 Zhang, Y.-N. *et al. Single-component self-assembling protein nanoparticles displaying stabilized prefusion-closed hemagglutinin trimers for influenza vaccine development* (Cold Spring Harbor Laboratory, 2025).

13 Powell, A. E. *et al.* A Single Immunization with Spike-Functionalized Ferritin Vaccines Elicits Neutralizing Antibody Responses against SARS-CoV-2 in Mice. *ACS Cent Sci* **7**, 183-199 (2021). https://doi.org/10.1021/acscentsci.0c01405

14 Xu, D. *et al.* Design of universal Ebola virus vaccine candidates via immunofocusing. *Proceedings of the National Academy of Sciences* **121** (2024). https://doi.org/10.1073/pnas.2316960121

15 Dickey, T. H. *et al.* Design of a stabilized RBD enables potently neutralizing SARS-CoV-2 single-component nanoparticle vaccines. *Cell Reports* **42**, 112266 (2023). https://doi.org/10.1016/j.celrep.2023.112266

16 Konrath, K. M. *et al.* Nucleic acid delivery of immune-focused SARS-CoV-2 nanoparticles drives rapid and potent immunogenicity capable of single-dose protection. *Cell Reports* **38**, 110318 (2022). https://doi.org/10.1016/j.celrep.2022.110318

17 Corbett, K. S. *et al.* Design of Nanoparticulate Group 2 Influenza Virus Hemagglutinin Stem Antigens That Activate Unmutated Ancestor B Cell Receptors of Broadly Neutralizing Antibody Lineages. *mBio* **10** (2019). https://doi.org/10.1128/mbio.02810-18
18 Kanekiyo, M. *et al.* Rational Design of an Epstein-Barr Virus Vaccine Targeting the Receptor-Binding Site. *Cell* **162**, 1090-1100 (2015). https://doi.org/10.1016/j.cell.2015.07.043
19 Yassine, H. M. *et al.* Hemagglutinin-stem nanoparticles generate heterosubtypic influenza protection. *Nature Medicine* **21**, 1065-1070 (2015). https://doi.org/10.1038/nm.3927
20 Sliepen, K. *et al.* Induction of cross-neutralizing antibodies by a permuted hepatitis C virus glycoprotein nanoparticle vaccine candidate. *Nature Communications* **13** (2022). https://doi.org/10.1038/s41467-022-34961-8
21 Chao, C. W. *et al.* Protein nanoparticle vaccines induce potent neutralizing antibody responses against MERS-CoV. *Cell Reports* **43**, 115036 (2024). https://doi.org/10.1016/j.celrep.2024.115036
22 McLeod, B. *et al.* Vaccination with a structure-based stabilized version of malarial antigen Pfs48/45 elicits ultra-potent transmission-blocking antibody responses. *Immunity* **55**, 1680-1692.e1688 (2022). https://doi.org/10.1016/j.immuni.2022.07.015
23 Malhi, H. *et al.* Immunization with a self-assembling nanoparticle vaccine displaying EBV gH/gL protects humanized mice against lethal viral challenge. *Cell Reports Medicine* **3**, 100658 (2022). https://doi.org/10.1016/j.xcrm.2022.100658
24 Sun, C. *et al.* A gB nanoparticle vaccine elicits a protective neutralizing antibody response against EBV. *Cell Host & Microbe* **31**, 1882-1897.e1810 (2023). https://doi.org/10.1016/j.chom.2023.09.011
25 He, L. *et al.* Single-component multilayered self-assembling nanoparticles presenting rationally designed glycoprotein trimers as Ebola virus vaccines. *Nature Communications* **12** (2021). https://doi.org/10.1038/s41467-021-22867-w
26 Mu, Z. *et al.* mRNA-encoded HIV-1 Env trimer ferritin nanoparticles induce monoclonal antibodies that neutralize heterologous HIV-1 isolates in mice. *Cell reports* **38**, 110514-110514 (2022). https://doi.org/10.1016/j.celrep.2022.110514
